# Enhancing grapevine breeding efficiency through genomic prediction and selection index

**DOI:** 10.1101/2023.07.31.551371

**Authors:** Charlotte Brault, Vincent Segura, Maryline Roques, Pauline Lamblin, Virginie Bouckenooghe, Nathalie Pouzalgues, Constance Cunty, Matthieu Breil, Marina Frouin, Léa Garcin, Louise Camps, Marie-Agnès Ducasse, Charles Romieu, Gilles Masson, Sébastien Julliard, Timothée Flutre, Loïc Le Cunff

**Affiliations:** UMT Geno-Vigne^®^, IFV, INRAE, Institut Agro Montpellier, 34398, Montpellier, France; Institut Français de la vigne et du vin, 30240, Le Grau du Roi, France; UMR AGAP Institut, Univ Montpellier, CIRAD, INRAE, Institut Agro Montpellier, 34398, Montpellier, France; Centre du Rosé, 70, Av. du Président Wilson, 83550 Vidauban, France; Conservatoire du Vignoble Charentais, Trepsec, 16370 Cherves-Richemont, France; Université Paris-Saclay, INRAE, CNRS, AgroParisTech, GQE – Le Moulon, 91190, Gif-sur-Yvette, France

## Abstract

Grapevine (*Vitis vinifera*) breeding reaches a critical point. New cultivars are released every year with resistance to powdery and downy mildews. However, the traditional process remains time-consuming, taking 20 to 25 years, and demands the evaluation of new traits to enhance grapevine adaptation to climate change. Until now, the selection process has relied on phenotypic data and a limited number of molecular markers for simple genetic traits such as resistance to pathogens, without a clearly defined ideotype and was carried out on a large scale. To accelerate the breeding process and address these challenges, we investigated the use of genomic prediction, a methodology using molecular markers to predict genotypic values. In our study, we focused on two existing grapevine breeding programs: *Rosé* wine and *Cognac* production. In these programs, several families were created through crosses of emblematic and inter-specific resistant varieties to powdery and downy mildews. 30 traits were evaluated for each program, using two genomic prediction methods: GBLUP (Genomic Best Linear Unbiased Predictor) and LASSO (Least Absolute Shrinkage Selection Operator). The results revealed substantial variability in predictive abilities across traits, ranging from 0 to 0.9. These discrepancies could be attributed to factors such as trait heritability and trait characteristics. Moreover, we explored the potential of across-population genomic prediction by leveraging other grapevine populations as training sets. Integrating genomic prediction allowed us to identify superior individuals for each program, using multivariate selection index method. The ideotype for each breeding program was defined collaboratively with representatives from the wine-growing sector.

## Introduction

Plant breeding has been a key lever to adapt varieties to human use and the environment. The genetic gain obtained after one cycle of a breeding program is given through the breeder’s equation (Lush, 1937). It depends on the additive genetic variance of the population, the accuracy and intensity of selection, and the cycle length. In grapevine, this cycle length is about 20 to 25 years, when accounting for phenotyping new varieties (Töpfer and Trapp, 2022). Thus, grapevine breeding is critically long and hereafter the genetic gain is reduced. Because of its perennial nature, grapevine needs to be adapted to challenging conditions, in a increasingly variable environment, due to climate change (Santos et al., 2020).

In the past years, grapevine breeding in Europe has been focused on disease resistance to powdery and downy mildews (Eibach et al., 2007; Schneider et al., 2019; Töpfer and Trapp, 2022). The French INRAE-ResDur program generated a dozen of varieties, all with at least two major resistance genes for each disease. The whole selection process lasted around 15 to 20 years (Reynolds and TBX, 2015; Schneider et al., 2019). Thus, there is a critical need for accelerating this selection process, while accounting for other traits related to climate change. Marker-assisted selection (MAS) was used in the INRAE-ResDur program to early detect seedlings with all resistance genes. However, most quantitative traits involved in adaptation to climate change are under a complex genetic determinism, with possibly thousands of genes involved (Alonso-Blanco and Méndez-Vigo, 2014; Flutre et al., 2022). In that case, QTL detection results in many small effects often overestimated and that are not transferable through MAS to breeding (Beavis et al., 1994; Crossa et al., 2017; Meuwissen et al., 2016; Xu, 2003).

Genomic selection (GS) has been proposed to avoid these limitations, thanks to the availability of genome-wide markers (Bernardo, 1994; Meuwissen et al., 2001). In GS, all markers are analyzed together and their associated effects on the phenotypes are jointly estimated in a training set population (TS). Then, these effects are applied in a validation set population (VS), on which only genotypes are available (Heffner et al., 2009). GS has been widely applied to animal and plant breeding, with some scarce examples of applications in grapevine (Brault et al., 2021, 2022b; Flutre et al., 2022; Fodor et al., 2014; Migicovsky et al., 2017; Viana et al., 2016). Especially, GS has only been applied in a research context, with varieties not intended for breeding. GS allows to save time in the breeding programs but it offers other interests (Consortium et al., 2021). Indeed, using GS allows testing of more crosses and offspring because no phenotyping is needed. Then, the selection intensity is increased, as more genotypes are tested, increasing the selection gain according to the breeder’s equation. Concerning the selection accuracy, the impact of GS is balanced. On the one hand, GS implies concentrating phenotyping on the training population, with possibly more replications that can increase the heritability and accuracy of the model. On the other hand, using a GS model trained in a population genetically far from the selection population would reduce the predictive ability (Brault et al., 2022b). One challenge of GS is then to find a trade-off between the advantages and drawbacks of GS in terms of prediction accuracy.

Once predicted or observed genotypic values are acquired, the breeder needs to select the best individuals in the population, by taking into consideration several traits and making compromises. This can be streamlined with a linear multi-trait selection index. The most famous selection index is the Smith-Hazel index (Smith, 1936). Since then, other algorithms have been developed to account for the multicollinearity between the traits (Olivoto and Nardino, 2021; Rocha et al., 2018). In grapevine, the ideotype (i.e., the criteria to combine all traits to get the best performing variety in each environment) is complex, because the wine is a transformed product and its quality relies on many variables (Reynolds and TBX, 2015; Töpfer and Trapp, 2022). Such an ideotype is likely to vary across wine regions. Specifically, the grapevine ideotype will include traits for which the genetic value must be maximized or minimized (directional selection) and traits for which an optimum value would be sought (stabilizing selection). Moreover, quality traits such as acids, sugars, anthocyanins, tannins, and volatile compounds interact with yield-related variables (Reynolds and TBX, 2015).

This article describes and proposes an application of GS to two breeding programs of grapevine varieties. These two breeding programs were compared, with a similar design of experiments but various traits and ideotypes. First, we fitted a mixed linear model for each experiment to extract genotypic values, then we applied genomic prediction within the training set to estimate predictive ability. Finally, we used multi-trait selection index to select the most promising individuals from predicted genotypic values.

## Material and methods

### Design of experiment

Two breeding programs were compared: the Martell breeding program, funded by Martell company which produces Cognac and conducted by the conservatory of the Charente vineyards, INRAE and IFV in France; and the EDGARR breeding programs, conducted by the *Center for Rosé*, INRAE, and IFV in France for producing Rosé wine. Both programs included crosses between varieties emblematic of the region and varieties with polygenic resistance to powdery and downy mildews (inter-specific hybrids). In both programs, after MAS, unselected individuals from the crosses were planted in a pot to constitute the TS, for genotyping and phenotyping while selected individuals were only genotyped and constitute the VS, except for a few families only present in the VS (Figure S1).

The Martell program included four famous grape varieties (Monbadon, Montils, Rayon d’Or, and Vidal 36), and the EDGARR experiment included two famous grape varieties (Cinsaut and Vermentino). The genetic relatedness between the individuals of the TS and the VS could be full-sibs, half-sibs, or no genetic relationship. A major difference between these programs was the number of genotypes. In the Martell program, there were 347 and 277 individuals in the TS and VS, respectively. In the EDGARR program, there were 193 and 132 individuals in the TS and VS, respectively.

### Genomic data analysis

The same genotyping approach was used in both programs. Genotyping was done using the genotyping-by-sequencing technology, using the *ApeKI* restriction enzyme (Elshire et al., 2011). Keygene N.V. owns patents and patent applications protecting its Sequence Based Genotyping technologies.

For EDGARR and Martell programs, SNP markers with more than 10% missing data and with less than 20 reads were discarded, producing 27,271 and 10,602 remaining SNPs, respectively. These markers were mapped to the updated version of the reference genome, PN40024.v4.2 (Velt et al., 2023). Genotypes with more than 50% missing data were also discarded. The remaining SNPs were imputed using Beagle software version 5.4 (Browning et al., 2018). Markers with a minor allele frequency lower than 1% were removed, giving a final table for EDGARR of 19,228 SNP markers for 326 individuals and for Martell of 10,380 SNPs for 624 individuals.

Then, for each cross, outlying individuals were detected using the Mahalanobis distance (Mahalanobis, 1936), with a p-value of 1%.

### Phenotypic data analysis

#### EDGARR

The individuals in the training population (e.g., 193 genotypes) were planted in pots, without rootstock. The vinestocks were managed to fast fruiting to accelerate the production of grapes. The experimental trial was located at the Espiguette domain, in Grau-du-Roi, in the South of France (43°29’48.5”N 4°08’13.2” E). There was one repetition per genotype, except for a small number of repeated controls (Cinsaut, Vermentino, Grenache and Syrah, with 5 or 6 repetitions).

In this population, 30 traits were phenotyped for two years (2018 and 2019), and 5 additional traits were phenotyped for one year. Traits were divided into five categories, namely acids with cis- and trans-coutaric acids, caftaric, ascorbic, hydroxycinnamic, malic, shikimic, tartaric acids, pH, and total acidity; color traits with blue, yellow and red absorbance, lightness, yellow and red indices, color intensity, tint and polymeric pigments at 420 and 520 nm; sugar traits with glucose and fructose; polyphenol traits with total polyphenol index, anthocyanin concentration; and finally agronomic and technologic traits with berry weight, glutathione, number of clusters and harvest date. A full description of these traits and summary statistics can be found in table S2 and table S3. Clusters were sampled when the sugar content reached 22° brix (gram of saccharose / 100 g). Some traits were measured with two non-redundant units: in concentration (g/l) and in amount in berries (mg/g of berries).

For the extraction of genotypic values, we first applied a full mixed model for each trait phenotyped for two years:

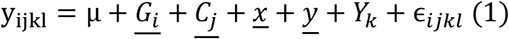, with *y*_*ijkl*_ the phenotypic observation for a given genotype i, cross j, year k and repetition l (for controls), *μ* the intercept, <u>*G*_*i*_</u> the random effect of the genotype j (nested in cross i), <u>*C*_*i*_</u> the random effect of the cross, <u>*x*</u> and <u>*y*</u> the random effects for coordinates of the plant in the trial, *Y*_*k*_ the fixed effect of the year (two levels), and ∈_*ijk*_ the residuals, assumed normally distributed. This full model was fitted with maximum likelihood, random effects were selected by a likelihood ratio test, and fixed effects were selected based on Fisher tests, using the lmerTest R-package (Bates et al., 2014). Variance components were estimated with restricted maximum likelihood on the selected model. The broad-sense heritability (H^2^) was computed as:

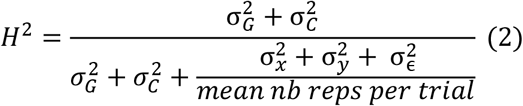

With 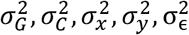 variances associated to genotype, cross, plot coordinates and residuals. Fitting information for all traits is available in table S4. If the genotype effect was not selected in the model, we re-fitted a simpler model with only the genotype effect as a random effect and we applied model selection only for fixed effects.

Best Linear Unbiased Predictors (BLUPs) were computed as the sum of the genotypic and cross effects (when cross effect was selected). We deregressed the BLUPs with the following formula: 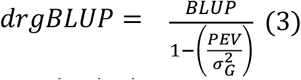 (Andrade et al., 2019; Garrick et al., 2009), with PEV the prediction error variance, i.e., the error associated with each BLUP value (for genotype and cross effects). This was estimated by the “postVar” parameter in ranef function from the lme4 R-package. For traits measured for one year, averaged phenotypic data per genotype was used.

#### Martell

For the Martell program, individuals were also planted in pots for the training population, without rootstocks, in Cognac region (45°44’22.9”N 0°21’58.2” E). The training set included 358 genotypes, among them, 349 came from progenies and 9 were grafted field controls (repeated 5 times). The phenotyping was done in 2021 and 2022 on potted plants for the training population. We studied 30 traits, which can be classified into 6 categories: vigor, disease, phenology, agronomic, technologic, and vinification. A full description of these traits and summary statistics can be found in table S2 and table S3. Traits related to harvest were sampled at around 10 alcohol content for the referent genotype (Ugni blanc).

The mixed model equation for phenotypic data analysis included effects described in (1) and some other effects: 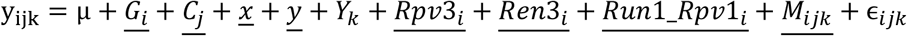, with as supplementary effects, the resistance genes Rpv3, Ren3 and Run1_Rpv1, and M a factor indicating the presence of available vine spur (if one spur and one cane were present, the pruning was simple guyot). We used the same equation (2) for computing the heritability for Martell population.

### Genomic prediction

The same pipeline of analysis was applied to both programs. First, genomic prediction (GP) was applied to the training population for all traits available with K-fold cross-validation, repeated R=10 times, with K=5. We implemented two genomic prediction methods: GBLUP with rrBLUP R package (Endelman, 2011) and the LASSO (Tibshirani, 1996), with glmnet R package (Friedman et al., 2010). GBLUP is more adapted to a complex genetic architecture (many QTLs), while LASSO is more adapted to a simpler genetic architecture. Predictive ability (PA) was estimated as Pearson’s correlation between observed and the predicted genotypic values. PA values were averaged across folds and cross-validation repetitions and standard errors were calculated.

The best method among the two was chosen for each population and trait and used to predict the genotypic values for the VS. The model was refit on the whole TS (without cross-validation) for predicting genotypic values for the VS. These values were deregressed using equation (3). For the LASSO, the deregressed values were obtained by fitting the Ordinary Least Square estimator for all selected markers in the training set.

For the EDGARR experiment, we predicted the berry color (red or white) using a logistic generalized linear model (GLM), adapted to binomial data with the LASSO method, using the glmnet R package, with as options family=‘binomial’ and alpha=1 (Friedman et al., 2010).

### Selection index

The selection index was designed by representatives of the wine growers for each of the two studied wine regions. It included traits for which the value needs to be maximized or minimized and traits for which an optimal value is required. The first selection criterion was the presence of the resistance genes for powdery and downy mildews, and the flower sex, handled with MAS.

The resulting multivariate selection index was computed using the MGIDI method (multi-trait genotype-ideotype distance index), described in (Olivoto and Nardino, 2021). Briefly, it rescales the phenotype on a 0-100 scale, in which 100 represents the maximum or the minimum value, depending on the direction of the selection. Then it performs a factor analysis, to summarize the multi-trait phenotypes and to avoid collinearity. Finally, the MGIDI is given by the sum of the distance between the actual phenotype and the ideotype for each factor. When an optimal value was sought by professionals, we computed the difference between the optimal value and the phenotype.

The selection index was applied for both programs, on predicted and deregressed genotypic values for the validation set individuals. The output of the MGIDI method included a strength and weakness view of selected individuals, with the contribution of each factor to the distance to the ideotype, and the rank of individuals, ordered by increasing MGIDI value.

### Other phenotypic and genomic data

We used genomic and phenotypic data from two other grapevine populations. A half-diallel population composed of 628 individuals from 10 bi-parental crosses where 5 parents were involved (Tello et al., 2019), phenotyped between 2013 and 2017. The second population is a diversity panel population of 277 genotypes, chosen to represent the genetic diversity of *Vitis vinifera* (Nicolas et al., 2016), and phenotyped between 2011 and 2012. Phenotypic and genomic data from these populations were already analyzed for genomic prediction and QTL detection in previous studies (Brault et al., 2022b, 2022a; Flutre et al., 2022).

There were 6 and 5 common traits with EDGARR and Martell programs, respectively. For genomic data, we performed a BLAST (Basic Local Alignment Research Tool) analysis on flanking sequences to find out the marker positions corresponding to the last version (PN40024.v4) of the *Vitis vinifera* reference genome (Velt et al., 2023). Then, we kept the common markers between each population and the target one (number of left SNPs displayed in Table S5). We fitted a GP model using GBLUP and LASSO for half-diallel, diversity panel, or both populations and kept the best method to predict genotypic values of EDGARR and Martell populations. We measured the predictive ability and compared it to the values from within-population GP.

## Results

### Genetic structure

For the EDGARR program, 325 individuals have been genotyped for 19,228 SNP markers after filters. For the Martell program, there were 624 individuals genotyped for 10,380 SNPs. A principal component analysis (PCA) was conducted to explore the genetic structure of the population. We found that families were well separated, located between their parents. Individuals in training and validation sets displayed a clear overlap, except for some families only in the validation set (Figure S1). The PCA analysis showed some outlier individuals, spotted with the Mahalanobis distance. For the EDGARR population, we excluded 3 individuals, all from Cinsaut x 3421-F02-PL5 cross; for the Martell population, we excluded 4 individuals from 4 crosses.

Overall, for both populations, the relative position of families seems to be driven by the inter-specific resistant parents. This is likely because they show more genetic diversity compared to *V. vinifera* varieties.

### Phenotypic structure

For both populations, we included environmental cofactors, despite a small number of repetitions. We were able to estimate the effects of plot position, year, and resistance gene, depending on the trait (Table S4). Broad-sense heritability values displayed a wide range across all traits. They ranged from 0 to 0.76 (average of 0.39) and from 0.002 to 0.99 (average of 0.43) for EDGARR and Martell populations, respectively. From the BLUPs of genotypic values, we applied a deregression to retrieve the original scale of the data in terms of mean and variance. We checked visually the quality of deregression. The correlation between raw averaged phenotypic data and deregressed BLUPs was between 0.73 and 0.98 for the EDGARR population, and between 0.60 and 0.99 for the Martell population. The matrix of pairwise genotypic correlations between the traits showed for EDGARR, that traits related to color were correlated to each other. Overall, for the other traits, genotypic correlations were mostly low (data not shown).

The PCA analysis showed a mild phenotypic structure (Figure S6). For EDGARR, the structure was driven by the inter-specific parents (F02-PL5 and F10-PL2) and by traits related to color, while for Martell, the crosses were more separated from each other, and the differentiation was driven by acid and yield traits.

### Genomic prediction results

Predictive abilities were comparable for both populations and covered a wide range of values between 0.04 to 0.87 (Figure 3). To avoid the effect of the genetic architecture on the predictive ability, we chose the best method between GBLUP and LASSO. Overall, GBLUP provided a better PA than LASSO for both populations, with an average of 0.41 and 0.34 for EDGARR and 0.44 and 0.39 for Martell, for GBLUP and LASSO, respectively. For EDGARR, GBLUP yielded a higher PA than LASSO for 26 traits out of 35, and 28 out of 30 for Martell. PA and heritability values were correlated for both populations, with a correlation value of 0.60 for EDGARR, and 0.42 for Martell. The different trait categories were quite evenly represented across the range of PA for both populations (Figure 3). However, traits for which the cross effect was not kept in the mixed model (1), displayed a lower PA with an average difference of 0.55 and 0.37 for EDGARR and Martell populations, respectively (Figure S7). We found that the 4 traits measured on a semi-quantitative scale for Martell populations had a slightly lower PA (difference of 0.26, a p-value of 0.044 using a Wilcoxon test). For EDGARR data, we could not fit a mixed model for 6 traits, because they were phenotyped in a single year. For these traits, averaged phenotypic data per genotype were used for GP. We found an average PA of 0.17 for these traits, up to 0.51 for trans-coutaric acid and the GBLUP method.

The predicted genotypic values were deregressed a second time in order to retrieve the initial mean and variance for each trait for applying our selection index. We computed the correlation between the genotypic values and the predicted values and visually checked that the scales were comparable. The correlations ranged from 0.42 to 1 (average of 0.82) for the EDGARR population, and from 0.24 (for vigor trait) to 1 (average of 0.87) for the Martell population.

We used a GLM with the LASSO method to predict categorical color for EDGARR population. The accuracies ranged between 0.943 to 0.963, with an average of 0.952 in cross-validation.

### Selection index

For both programs, the selection index was established by the professional committee in charge of local grapevine breeding. The first criterion was the presence of two resistance genes both to powdery and downy mildews. Then, a specific index was determined, based on the traits available.

### EDGARR selection index

For EDGARR population, the center of *Rosé* established a selection index to get varieties with more acidity, less color, higher productivity, and adaptation to climate change. Finally, the corresponding ideotype included 11 traits, 5 traits to be optimized (must tartaric, malic, total acidity, pH, alcohol content), 2 traits to be minimized (color intensity and total polyphenol index) and 4 traits to be maximized (berry tint, number of clusters, berry weight and harvest date) (Table S8). We used PA values as weights associated with each trait. The MGIDI algorithm selected 3 factors, represented by 4 (tartaric acid, berry lightness, berry color, color intensity, harvest date), 3 (malic acid, total acidity, pH), and 4 traits (alcohol content, total polyphenol index, number of cluster and berry weight), respectively (Table S9). Vermentino was a parent of 12 out of 15 selected genotypes, and 8 individuals from the same cross Vermentino x F10-PL2 were selected. Surprisingly, the resistant genotype F02-PL5 was not selected as a parent of the first 15 genotypes. From the PCA analysis (Figure 4), it is clear that selected individuals are phenotypically close to each other. The predicted berry color was white for 4 genotypes, and the genotype with the lowest MGIDI was predicted white. Factors 1 and 2 contributed the most to the MGIDI score for the selected genotypes, which means that they performed quite similarly for factor 3. Some genotypes performed better for some factors, such as P869-F04 for factor 1, or P596-A09 for factor 2, while others had a more balanced performance across factors, such as P249-F10 (Figure S12, Table S9).

### Martell selection index

The ideotype for Martell included 11 traits, 7 traits to be maximized (global and primary fertility, yield, tartaric acid, total must acidity, cluster weight and berry weight), 1 to be minimized (must pH), and 2 with an optimum value (must malic acid and ease of detachment of pedicel, OIV 240) (Table S8). We excluded beforehand traits with a PA value lower than 0.5. The MGIDI algorithm selected 3 factors, represented by 5 (total acidity, pH, yield, primary and global fertility), 4 (cluster weight, tartaric acid, berry weight, malic acid) and 2 traits (ease of detachment of pedicel and potassium), respectively (Table S9). Among the selected traits, some of them displayed high genetic correlations (positive or negative). The average of the 15 genotypes selected followed the expected trend (increase or decrease compared to the average of the population), for all the traits, except for single berry and cluster weights (Table S9). Distributions of predicted genotypic values and the position of some parents and selected genotypes are displayed in Figure S10. For 13 out of 15 genotypes selected, Monbadon and C03-PL5 were one of the two parents (Table S11). As for EDGARR, factor 3 contributed less to the MGIDI score and genotypes displayed various strengths or weaknesses for the factors. In particular, the superior performance of E12-32G10 (ranked 1^st^) was due to factor 1 and 3, and E10-29D10 (ranked 7^th^) was only due to factors 2 and 3 (Figure S12, Table S9).

### Across-population genomic prediction

For EDGARR and Martell, within-population GP was better for 4 traits out of 6, and for 5 traits out of 5, respectively. PA values for across-population GP were variable, mostly depending on the trait, on the validation population, and to a lesser extent on the training population (Figure 5). Overall, across-population PA values were much higher in EDGARR than in the Martell population. For EDGARR and two traits (shikimic acid concentration and number of clusters), using data from the diversity panel and the half-diallel led to a higher PA than using data from the same population.

For EDGARR, using both data from the diversity panel and the half-diallel led to higher PA, except for the harvest date, for which a strong decrease was observed. For Martell, the diversity panel was the best training population consistently for all traits.

We used for each trait and training set, the best method among GBLUP and LASSO. The results show that GBLUP was the only method selected for within-population, while LASSO was the best method for at least one trait for both populations.

## Discussion

Our study comprised the analysis of 30 traits for two grapevine populations. Some of the individuals were only genotyped, which allowed us to perform genomic predictions. We first tested the ability of GP models to accurately predict the genotypic values in a within-population scenario. Then, we proposed a selection index and selected the most relevant individuals according to it. The ideotype was built in partnership with professional wine growers and was specific to each of the two wine regions studied. To our knowledge, this is the first time a precise ideotype is described for grapevine. Finally, some of the phenotyped traits were also available for other grapevine populations. We tested to train the GP model with these less related individuals for the common traits and the results were encouraging in one of the two populations.

### Comparison of the populations

The two populations studied were similar in the sense that they were composed of bi-parental crosses with a resistant and emblematic grapevine variety as parents (Figure 1). In both designs, the number of individuals per cross was highly unbalanced, especially in the validation set (Figure S1). We observed that number of remaining SNP markers was higher in EDGARR than in Martell population (27,271 and 10,602, respectively), despite a higher number of reads per genotype for Martell (4.6 M) compared to EDGARR (4 M). This might be explained by the broader genetic diversity and less genetic relatedness of the resistant parents in the Martell population.

**Figure 1.**
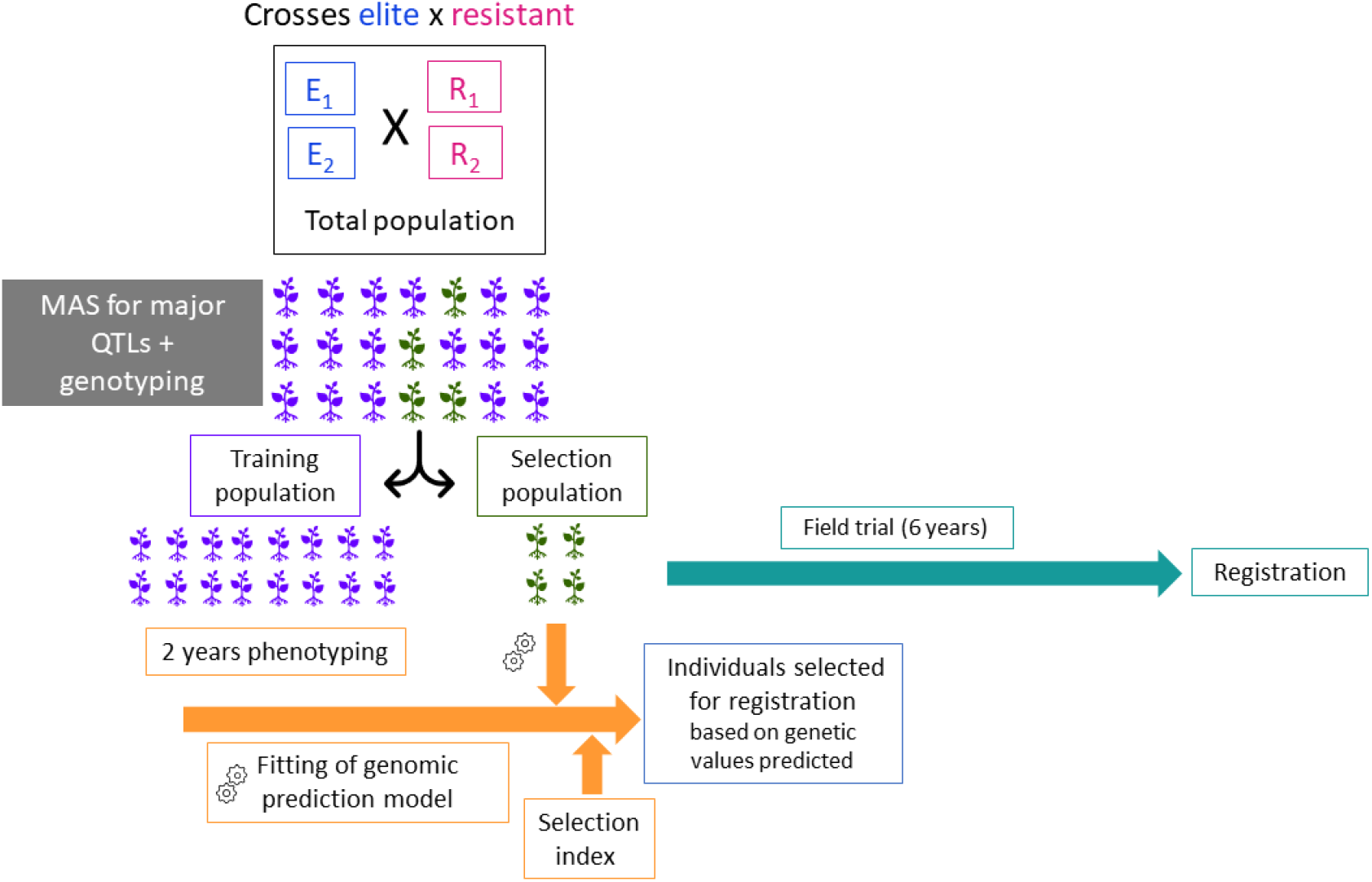
Design of experiment for EDGARR and Martell breeding programs.

**Figure 2.**
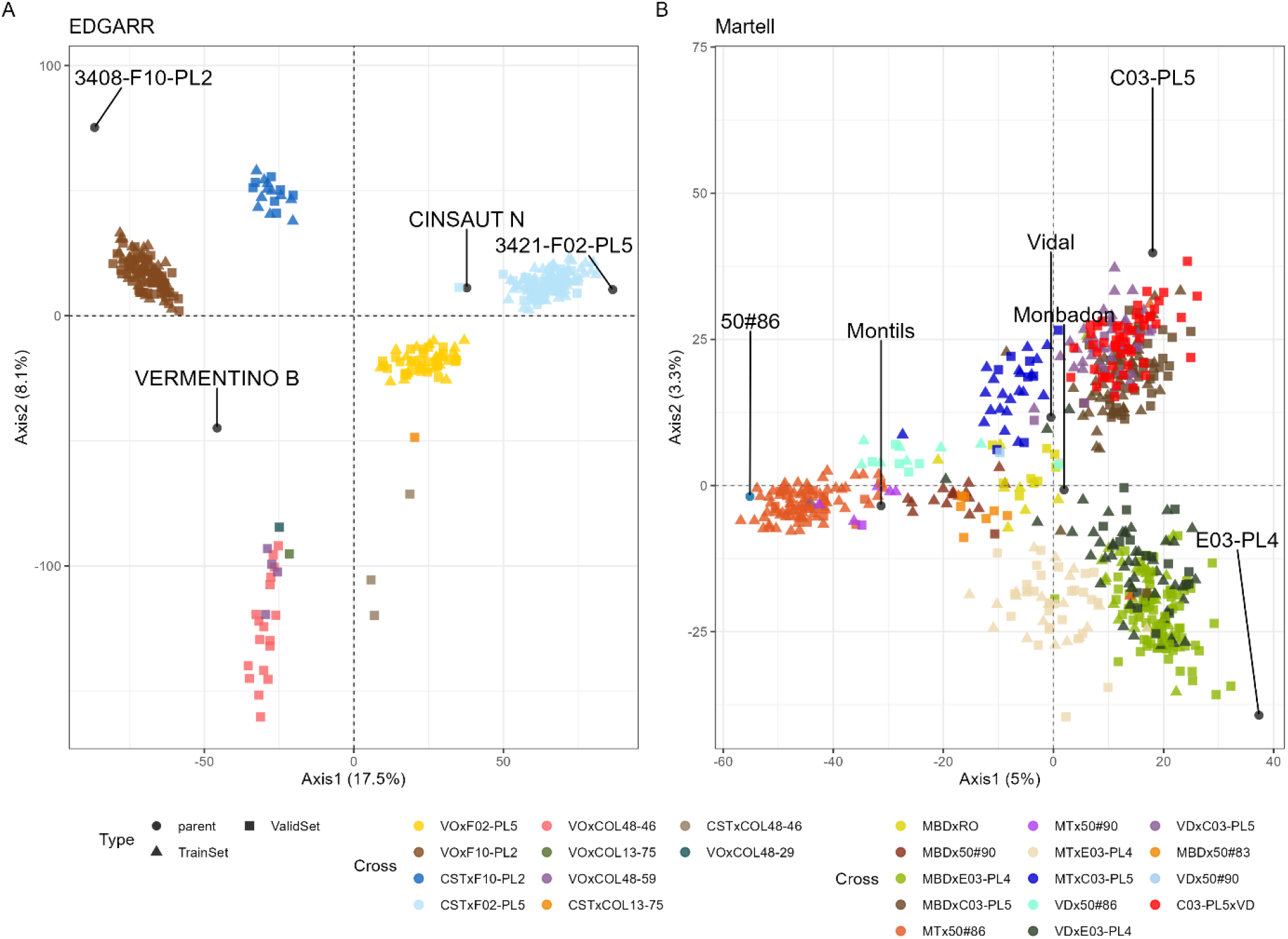
PCA of genetic markers for EDGARR (A) and Martell (B) populations. Parents are labelled. The point shape corresponds to the type of individual, triangle: training set, square: validation set, points: parents of crosses. Cross names were abbreviated as follow: Vermentino (VO), Cinsaut (CST), Monbadon (MBD), Rayon d’Or (RO), Montils (MT) and Vidal (VD).

**Figure 3.**
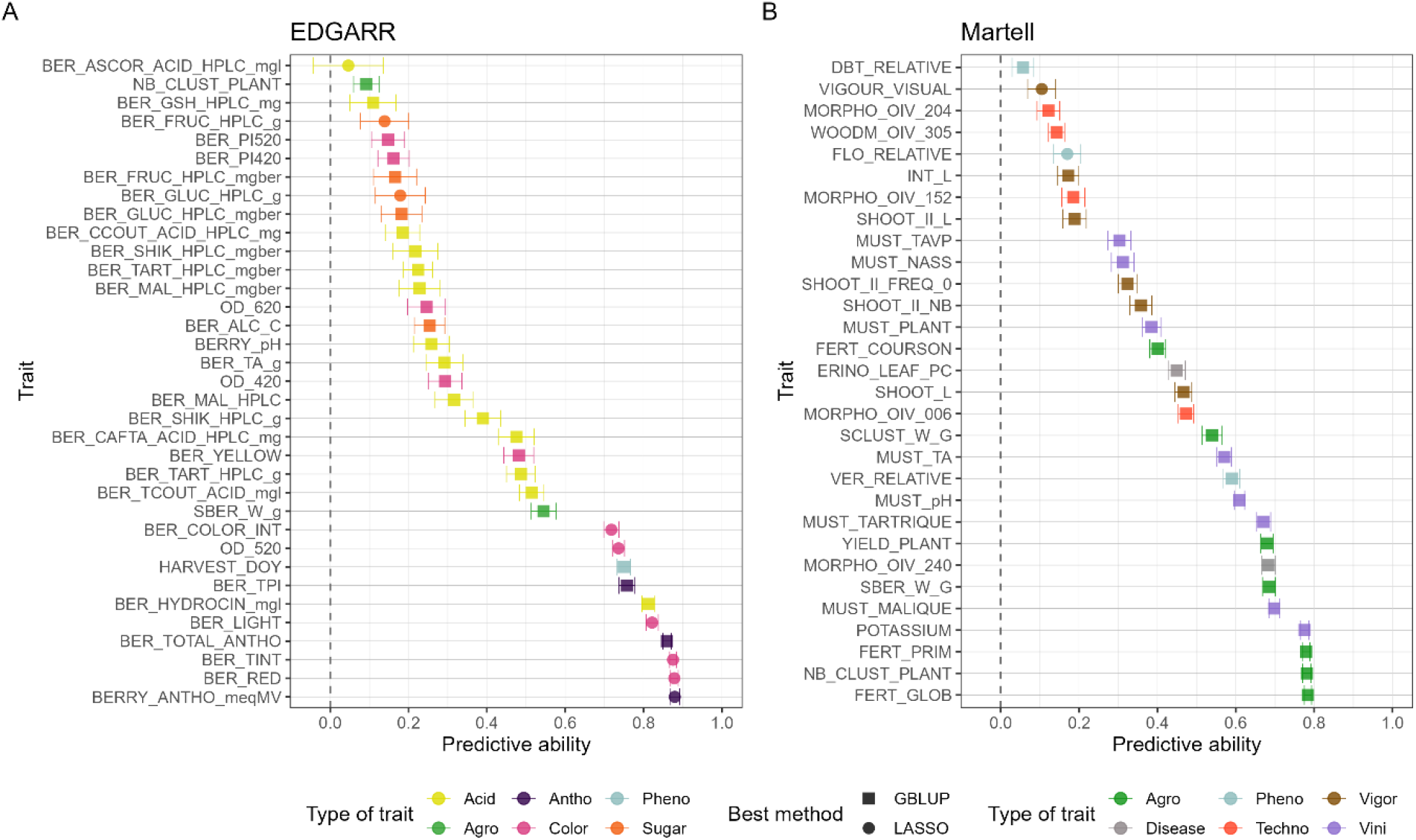
Predictive ability for all traits for EDGARR (A) and Martell (B) populations. Error bars correspond to standard errors calculated across cross-validation repetitions. For each trait, the best method among GBLUP and LASSO was selected.

**Figure 4.**
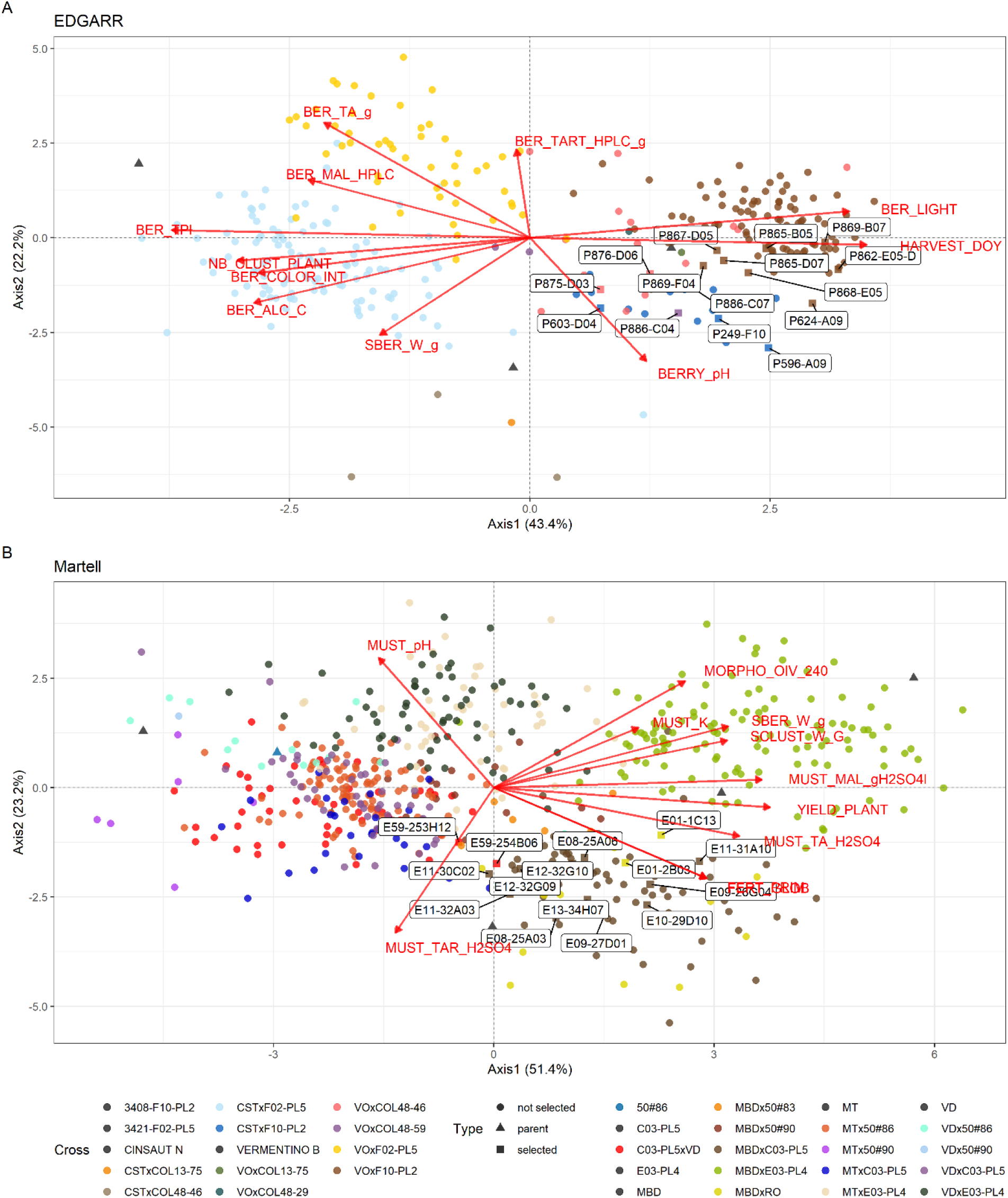
PCA of the genotypic values for the selection candidates for the traits in the selection index for the first two principal components. Variables are displayed in red, and genotypes are colored according to their cross. Selected individuals are labelled. A: EDGARR population, B: Martell population.

**Figure 5.**
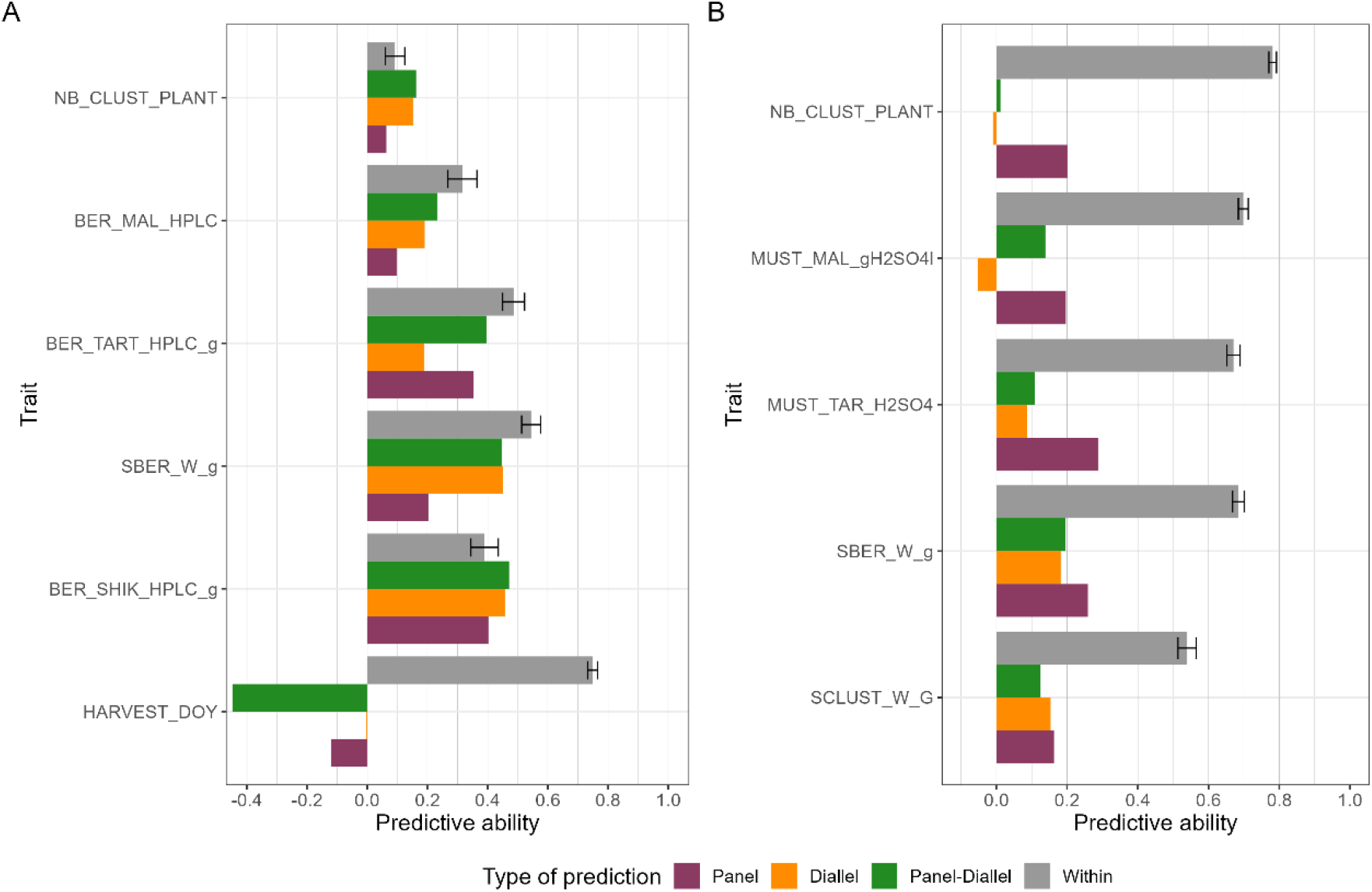
Comparison of the predictive ability for various training sets. A: EDGARR population, B: Martell population.

The size of the entire population for Martell was about twice the size of EDGARR. Nevertheless, PA observed were similar for both populations, with a comparable range and average. There were only two common traits between these populations: berry weight (SBER_W_g) and number of clusters (NB_CLUST_PLANT). Other traits were close, such as malic and tartaric acids, or total acidity, but they were not measured on the same entity (berry for EDGARR and must for Martell). Then, they were considered as different traits. PA for the number of clusters was extremely different in the two populations, with low PA for EDGARR (0.09), and high PA for Martell (0.78) (Figure 3). This might be explained by the fact that for EDGARR, the number and length of shoots were not controlled. Then, the number of clusters is relative to the number of shoots and the fertility. This result is consistent with the difference in heritability values between these populations (Figure S4). For EDGARR, traits for sugar concentrations displayed low heritability and PA, probably because the sampling date was determined by a sugar threshold, thus the genetic variability for these traits was minimized. These results illustrate the effect of vineyard management and measurement methodology on heritability values.

### Factors affecting the predictive ability

A major factor impacting PA was the presence of the cross effect in the final BLUP model (Figure S7). We found that traits with the cross effect had more differentiated genetic values per cross. Then, PA was automatically increased because we predicted both the average of a cross and the Mendelian sampling part (within a cross) (Werner et al., 2020; Würschum et al., 2017). This effect was highlighted by Werner et al., (2020), who measured PA per cross and for several crosses. However, we could not use a single cross as training or validation population, because we did not have enough genotypes and cross sizes were unbalanced. Brault et al. (2022b), compiled predicted genotypic values per cross and calculated the PA of GP for each cross and PA for predicting the cross means. But again, we had too few individuals to accurately measure PA for each cross.

As expected, the heritability values were overall correlated with PA values for both populations.

Across-population GP was competitive with within-population GP for the EDGARR population (Figure 5). This was unexpected since the TS used in across-population scenario was phenotyped in the field and during different years compared to EDGARR population, phenotyped in pots. In the Martell population, PA values were higher in the within-population scenario, and differences between within and across-population GP was higher compared to EDGARR population. However, we observed that PA values in across-population were higher for EDGARR than for Martell for SBER_W_g (Figure 5), while TS sizes were constant. Then, this difference in PA in the across-population scenario could be due to the differences in genetic relatedness between TS and VS, or by the phenotyping environment. The diversity panel and the half-diallel were planted about 20 km apart from the EDGARR population, and about 400 km apart from the Martell population. Our results suggest that the geographic proximity of TS and VS could have more impact on PA than genetic relatedness or TS size.

In the Martell population, we studied semi-quantitative traits, which displayed slightly smaller PA than other traits (Figure S7). We considered such traits as normal traits, even if the assumption of normality was strongly violated. Recently, (Azevedo et al., 2023) showed that using a linear mixed model for GP of ordinal traits was robust but sub-optimal. They advised using Bayesian Ordinal Regression Models, even though it is computationally demanding.

### Future breeding programs

These breeding programs aimed to save time and maximize the genetic relatedness between training and validation sets. First, individuals were filtered by MAS for disease resistance and hermaphroditism (Figure 1). The discarded individuals were quickly planted in pots to be phenotyped and serve as the TS, while genotypic values could be predicted for the VS, using GP. Such a breeding program relies on two strong hypotheses: i) phenotypes do not display a high genotype-by-environment (GxE) interaction between pots and the field, and ii) genetic relatedness is a major parameter of PA. Indeed, if we observe a strong GxE interaction, the ranking of individuals between pots and field will likely vary, hampering an accurate selection of the best individuals. To some extent, this was tested in the across-population scenario and PA values were nearly as high as they were in the within-population scenario for some traits for EDGARR population. This hypothesis should be further investigated for more traits and scenarios. For the second hypothesis, if genetic relatedness was already known to affect PA, its magnitude remains unknown. Especially in this study where the VS was composed of inter-specific varieties, while phenotypic data were only available for *Vitis vinifera* varieties. This is the first time that

GP has been applied with such different genetic backgrounds between the training and the validation sets. We tested using completely different populations to train the model, and results were encouraging for most traits for EDGARR population, while PA values were smaller in across-population for Martell population.

For across-population GP, we showed that LASSO was more often better than GBLUP, compared to the within-population scenario. This observation was also done in another study on grapevine (Brault et al., 2022b).

### Phenotyping environment

In our design of experiment, there was no repetition of a given genotype for a given year. Despite this, we could have medium to high heritability values depending on the trait. These values must be taken with caution, as variance components are likely not well estimated with this design.

Potted own-rooted grapevine phenotypes are likely to differ compared to field phenotypes. However, we have not found studies that compared both different varieties and traits related to the harvest. Most studies on pots or greenhouses were focused on disease resistance or drought tolerance. If this kind of breeding is chosen for the future, one should measure the GxE interactions beforehand.

### Grapevine ideotype

For EDGARR, a variety for *Rosé* wine was sought, with a little color, while for Martell, a variety for *Cognac* production was sought, with a white berry color, and high yield. Beyond those criteria, both projects were aiming to counter-balanced the effects of climate change on berry composition, namely higher alcohol degree, lower acidity, and shorter growth period (Bécart et al., 2022; Cortázar-Atauri et al., 2017; Parker et al., 2020; Rienth et al., 2021, 2016). These traits interact with each other’s. Selecting varieties that are ripening later (i.e., at the beginning of autumn in the Northern hemisphere) will experience lower temperatures during ripening, which would slow the degradation of malic acid and the accumulation of sugar (van Leeuwen et al., 2019). Ideotypes are now integrating traits related to the wine product, climate change, disease resistance, and more generally to production (yield, ability to produce wine). Other traits not directly in the ideotype would also be important, such as the resistance to black-rot *Guignardia bidwellii*, to *millerandage* and to *coulure* (poor fruit set). Besides, one may want to select individuals with medium performance across the traits or to correct the default of current grape varieties. The last solution is possible only if musts are blended.

As many other traits could not be included in the ideotype because of the difficulty of phenotyping, one must ensure that the selection intensity is not too high. Thus, enough individuals with genetic diversity must be kept to be phenotyped for costly traits such as wine aromas later in the breeding program.

Another solution for grapevine breeding would be to predict the best crosses to realize, based on the cross mean and variance prediction. The proof-of-concept for cross mean was already done in grapevine (Brault et al., 2022b) but it was not applied in a breeding context. Predicting cross variance would allow to select crosses that would result in extreme offspring phenotypes (Neyhart and Smith, 2019; Wolfe et al., 2021).

In contrast to other crops, the grapevine ideotype is likely to include traits for which an optimum value is sought. That is why we used deregressed genetic values so that the range of values for these traits remains meaningful to breeders. However, such double deregression as we did here could hamper the prediction quality. For the mixed model, we could have used BLUEs instead of BLUPs, but the design of experiment was too unbalanced, especially for the number of individuals per cross.

## Conclusion

This study provided the first insights on how genomic prediction could be integrated into grapevine breeding programs. The comparison of two breeding programs helped us identify factors affecting the prediction accuracy and determining the best conditions for applying genomic prediction, notably the training population environment and phenotypic reliability. For the first time in grapevine, a multi-trait selection index was used based on predicted genotypic values to help select the best cultivars.

## Data availability

Data and code to reproduce the results are available at: https://doi.org/10.57745/G8PXEJ.

Genomic and phenotypic data for half-diallel and diversity panel populations are available at: https://doi.org/10.15454/PNQQUQ.

## Contribution statement

C.B performed data curation, statistical analysis and wrote the manuscript with inputs from all authors; V.S and T.F contributed to the design, V.S, L.LC and T.F aided in interpreting the results and worked on the manuscript; L.LC designed and directed the project; M.R performed genomic experiments and bioinformatics analysis; P.L, V.B, M.B, M.F, L.G, L.C, M-A.D, C.R and L.LC carried out phenotyping and data management; L.LC, N.P, C.C and G.M were in charge of the EDGARR program; L.LC, M.F and S.J in charge of the Martell program.

