## Supplement materials for "Enhancing grapevine breeding efficiency through genomic prediction and selection index"

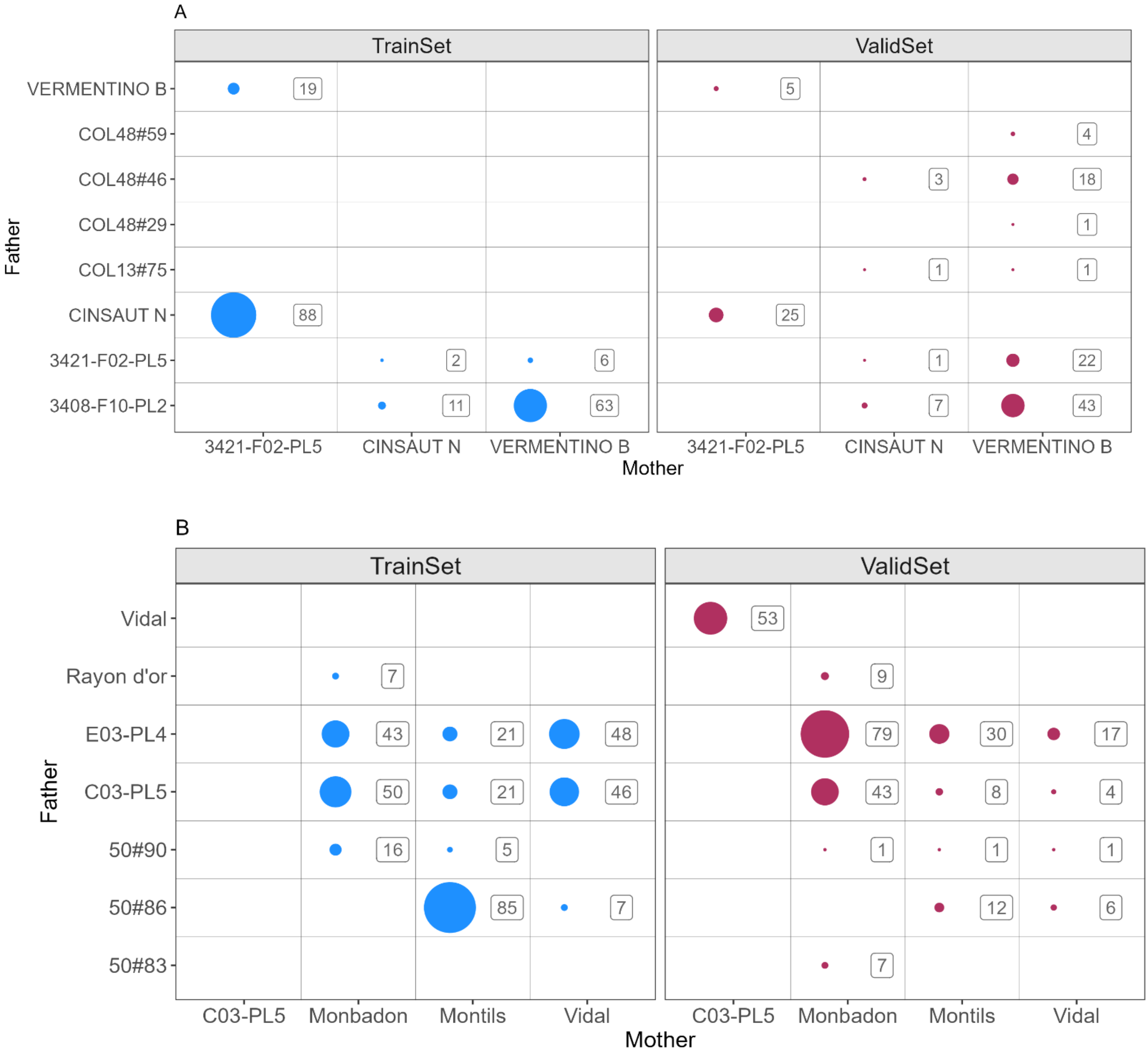

Figure S1 Number of offspring for each family depending on the category (training set and validation set). A: EDGARR, B: Martell.

| EDGARR | | | |
|---|---|---|---|
| Variable | description | VitisOntology_corresp | category |
| BERRY_ANTHO_meqMV | Berry Anthocyanins content (in meq of malvidin) | NA | Antho |
| BERRY_pH | Measure of pH on berries | CO_356:1000184 | Acid |
| BER_ALC_C | Potential alcohol content in berries | NA | Sugar |
| BER_ASCOR_ACID_HPLC_mgl | Concentration of ascorbic acid in berries, measured by HPLC in mg/L | NA | Acid |
| BER_CAFTA_ACID_HPLC_mg | Concentration of caftaric acid in berries, measured by HPLC in mg/L | NA | Acid |
| BER_CCOUT_ACID_HPLC_mg | Concentration of cis-coutaric acid in berries, measured by HPLC in mg/L | NA | Acid |
| BER_COLOR_INT | Color intensity measured on berries, determined by the sum of the absorbances at 420, 520 and 620 nm | NA | Color |
| BER_FRUC_HPLC_g | Fructose concentration in the berries (HPLC, g/l) | NA | Sugar |
| BER_FRUC_HPLC_mgber | Fructose amount per berry (HPLC, mg/berry) | NA | Sugar |
| BER_GLUC_HPLC_g | Glucose concentration in the berries (HPLC, g/l) | NA | Sugar |
| BER_GLUC_HPLC_mgber | Glucose amount per berry (HPLC, mg/berry) | NA | Sugar |
| BER_GSH_HPLC_mg | Glutathione concentration in the berries (HPLC, mg/l) | NA | Acid |
| BER_HYDROCIN_mgl | Concentration of hydroxycinnamic acid in berries, measured by the absorbance at 320 nm, in mg/l. | NA | Acid |
| BER_LIGHT | Perceptual lightness of berries, 0 to 100 for darkness to white (L* parameter). | NA | Color |
| BER_MAL_HPLC | Malic acid concentration in the berries (HPLC, g/l) | CO_356:1000299 | Acid |
| BER_MAL_HPLC_mgber | Malic acid amount per berry (HPLC, mg/berry) | NA | Acid |
| BER_PI420 | Polymeric pigments measured on berries by the absorbance at 420 nm | NA | Color |
| BER_PI520 | Polymeric pigments measured on berries by the absorbance at 520 nm | NA | Color |
| BER_RED | Redness measured on berries on a green to red scale (a* parameter). | NA | Color |
| BER_SHIK_HPLC_g | Shikimic acid concentration in the berries (HPLC, g/l) | NA | Acid |
| BER_SHIK_HPLC_mgber | Shikimic acid amount per berry (HPLC, mg/berry) | NA | Acid |
| BER_TART_HPLC_g | Tartaric acid concentration in the berries (HPLC, g/l) | CO_356:1000301 | Acid |
| BER_TART_HPLC_mgber | Tartaric acid amount per berry (HPLC, mg/berry) | NA | Acid |
| BER_TA_g | Titratable acidity of the berries (g/l H2SO4) | CO_356:1000302 | Acid |
| BER_TCOUT_ACID_mgl | Concentration of trans-coutaric acid on berries, measured by HPLC in mg/L | NA | Acid |
| BER_TINT | Tint or hue measured on berries, corresponding of the ration between the 420 and 520 nm absorbances. | NA | Color |
| BER_TOTAL_ANTHO | Total anthocyans measured on berries, by the absorbance at 520 nm | NA | Antho |
| BER_TPI | Total polyphenol index measured in berries, determined by the absorbance at 280 nm. | NA | Antho |
| BER_YELLOW | Yellowness measured on berries on a blue to yellow scale (b* parameter). | NA | Color |
| HARVEST_DOY | Harvest date (Day of year) | CO_356:1000289 | Pheno |
| NB_CLUST_PLANT | Number of clusters per plant | CO_356:1000158 | Agro |
| OD_420 | Optical density at 420 nm (yellow color) | NA | Color |
| OD_520 | Optical density at 520 nm (red color) | NA | Color |
| OD_620 | Optical density at 620 nm (blue color) | NA | Color |
| SBER_W_g | Single berry weight | CO_356:2000144 | Agro |

| Martell | | | |
|---|---|---|---|
| Variable | Description | VitisOntology_corresp | category |
| BUD_RELATIVE_CHAS | Bud burst date, relative to the mean bud burst date of Chasselas from plot | NA | Pheno |
| FERT_GLOB | Number of inflorescences per shoot (all kinds of shoots) | https://cropontology.org/term/CO_356:1000266 | Agro |
| FERT_PRIM | Total number of inflorescences/number of primary shoots | https://cropontology.org/term/CO_356:1000267 | Agro |
| FERT_SPURS | Number of inflorescences per shoot on spurs | https://cropontology.org/term/CO_356:1000265 | Agro |
| FLO_RELATIVE_CHAS | Flowering date, relative to the mean flowering date of Chasselas from plot | NA | Pheno |
| INT_L | Internode length | https://cropontology.org/term/CO_356:1000093 | Vigor |
| INT_L_45 | Internode length of rank 4 and 5 in cm | NA | Vigor |
| INT_NB | Number of internodes | NA | Vigor |
| LEAF_ERINOSE_PC | Percentage of erinose symptoms on leaves | NA | Disease |
| MORPHO_OIV_006 | Shoot attitude | https://cropontology.org/term/CO_356:1000198 | Techno |
| MORPHO_OIV_152 | Insertion of 1st inflorescence | https://cropontology.org/term/CO_356:1000091 | Techno |
| MORPHO_OIV_204 | Bunch density | https://cropontology.org/term/CO_356:1000037 | Techno |
| MORPHO_OIV_240 | Berry: ease of detachment from pedice | https://cropontology.org/term/CO_356:1000018 | Techno |
| MUST_ALC_C | Probable alcohol content in must, in % | NA | Vini |
| MUST_K | Must Potassium concentration | https://cropontology.org/term/CO_356:1000188 | Vini |
| MUST_MAL_gH2SO4l | Concentration of malic acid in must (equivalence g/l of H2SO4) | NA | Vini |
| MUST_N_AA&NH4 | Nitrohen content of the must (sum of N-NH4 and N-amino ac.) in g/l | https://cropontology.org/term/CO_356:1000355 | Vini |
| MUST_TAR_H2SO4 | Concentration of tartaric acid in must, expressed in equivalence of H2SO4 in gl | NA | Vini |
| MUST_TA_H2SO4 | Must titratable acidity (equivalence g/l of H2SO4) | https://cropontology.org/term/CO_356:1000293 | Vini |
| MUST_Y_PLANT | Must yield per plant, in mL | NA | Vini |
| MUST_pH | Must pH | https://cropontology.org/term/CO_356:1000185 | Vini |
| NB_CLUST_PLANT | Number of clusters per plant | https://cropontology.org/term/CO_356:1000158 | Agro |
| SBER_W_g | Single berry weight in gram | https://cropontology.org/term/CO_356:1000215 | Agro |
| SCLUST_W_G | Cluster weight | https://cropontology.org/term/CO_356:1000048 | Agro |
| SHOOT_II_FREQ_0 | Frequence of shoots with no internodes | NA | Vigor |
| SHOOT_L | Primary shoot length | https://cropontology.org/term/CO_356:1000211 | Vigor |
| VER_RELATIVE_CHAS | Véraison date, relative to the mean véraison date of Chasselas from plot | NA | Pheno |
| VIGOUR_VISUAL | Visual vigour observation in a 1-9 scale (by 2) | NA | Vigor |
| WOODM_OIV_305 | Time of beginning of wood maturity | https://cropontology.org/term/CO_356:1000228 | Techno |
| YIELD_PLANT | Total weight harvest per plant | https://cropontology.org/term/CO_356:1000247 | Agro |

Table S2 Description of traits, variable name, definition, link to crop ontology.

| EDGARR | | | | | | | |
|---|---|---|---|---|---|---|---|
| Variable | N | Mean | Std. Dev. | Min | Pctl. 25 | Pctl. 75 | Max |
| HARVEST_DOY | 403 | 263.29 | 24.20 | 173.00 | 248.00 | 280.00 | 301.00 |
| NB_CLUST_PLANT | 343 | 10.81 | 7.59 | 0.00 | 5.00 | 15.00 | 41.00 |
| BER_GSH_HPLC_mg | 93 | 16.26 | 19.09 | 0.83 | 3.22 | 22.29 | 95.80 |
| BER_GLUC_HPLC_mgber | 179 | 340.54 | 79.65 | 174.90 | 286.76 | 376.54 | 708.15 |
| BER_GLUC_HPLC_g | 179 | 88.47 | 10.31 | 45.27 | 82.16 | 94.27 | 129.36 |
| BER_FRUC_HPLC_mgber | 179 | 337.95 | 78.77 | 169.33 | 282.66 | 372.51 | 659.49 |
| BER_FRUC_HPLC_g | 179 | 87.83 | 10.83 | 43.83 | 81.44 | 94.94 | 126.54 |
| BER_TART_HPLC_mgber | 179 | 30.98 | 8.39 | 17.27 | 24.51 | 35.90 | 56.92 |
| BER_TART_HPLC_g | 179 | 8.03 | 1.62 | 4.74 | 6.72 | 9.24 | 13.02 |
| BER_MAL_HPLC_mgber | 179 | 51.98 | 18.59 | 23.71 | 38.11 | 62.73 | 116.48 |
| BER_MAL_HPLC | 179 | 13.34 | 3.23 | 7.08 | 10.86 | 15.36 | 23.30 |
| BER_SHIK_HPLC_mgber | 178 | 0.10 | 0.10 | 0.00 | 0.03 | 0.13 | 0.75 |
| BER_SHIK_HPLC_g | 178 | 0.03 | 0.03 | 0.00 | 0.01 | 0.03 | 0.17 |
| BER_ASCOR_ACID_HPLC_mgl | 101 | 5.67 | 2.95 | 1.11 | 3.56 | 7.27 | 14.80 |
| BER_CAFTA_ACID_HPLC_mg | 101 | 142.56 | 59.85 | 38.88 | 102.19 | 176.63 | 312.28 |
| BER_TCOUT_ACID_mgl | 101 | 175.40 | 99.72 | 19.05 | 104.74 | 225.04 | 572.03 |
| BER_CCOUT_ACID_HPLC_mg | 101 | 33.94 | 18.54 | 7.24 | 18.93 | 42.58 | 100.95 |
| SBER_W_g | 267 | 390.25 | 125.47 | 46.54 | 308.36 | 467.96 | 789.90 |
| BER_ALC_C | 267 | 11.79 | 1.01 | 7.69 | 11.34 | 12.42 | 14.08 |
| BER_TA_g | 265 | 5.52 | 1.48 | 2.90 | 4.50 | 6.45 | 10.40 |
| BERRY_pH | 262 | 3.73 | 0.23 | 3.18 | 3.56 | 3.90 | 4.41 |
| BERRY_ANTHO_meqMV | 233 | 70.74 | 80.00 | -8.77 | 15.34 | 111.48 | 503.23 |
| BER_LIGHT | 272 | 63.21 | 17.61 | 0.06 | 50.09 | 77.08 | 95.42 |
| BER_RED | 272 | 28.31 | 18.48 | -2.04 | 13.95 | 42.52 | 66.00 |
| BER_YELLOW | 272 | 17.72 | 9.44 | -6.95 | 10.22 | 24.43 | 47.73 |
| OD_420 | 272 | 0.80 | 0.57 | 0.15 | 0.48 | 0.96 | 7.19 |
| OD_520 | 272 | 0.94 | 0.81 | 0.07 | 0.43 | 1.19 | 6.85 |
| OD_620 | 272 | 0.25 | 0.27 | 0.00 | 0.12 | 0.31 | 3.62 |
| BER_COLOR_INT | 272 | 1.99 | 1.56 | 0.28 | 1.04 | 2.50 | 17.66 |
| BER_TINT | 272 | 1.11 | 0.55 | 0.35 | 0.68 | 1.43 | 3.56 |
| BER_PI420 | 239 | 0.54 | 0.37 | 0.02 | 0.32 | 0.66 | 3.85 |
| BER_PI520 | 239 | 0.28 | 0.24 | 0.01 | 0.15 | 0.33 | 2.98 |
| BER_TPI | 235 | 13.11 | 5.11 | 4.61 | 9.64 | 15.60 | 43.65 |
| BER_HYDROCIN_mgl | 235 | 9.64 | 3.60 | 3.49 | 6.93 | 12.10 | 27.50 |
| BER_TOTAL_ANTHO | 235 | 3.99 | 4.10 | -0.27 | 1.21 | 5.89 | 26.21 |

| Martell | | | | | | | |
|---|---|---|---|---|---|---|---|
| Variable | N | Mean | Std. Dev. | Min | Pctl. 25 | Pctl. 75 | Max |
| BUD_RELATIVE_CHAS | 739 | 9.20 | 7.11 | -2.00 | 3.00 | 14.00 | 50.00 |
| FLO_RELATIVE_CHAS | 668 | 0.97 | 4.08 | -23.00 | -1.00 | 5.00 | 16.00 |
| VER_RELATIVE_CHAS | 661 | 22.34 | 10.49 | -10.00 | 11.00 | 35.00 | 45.00 |
| WOODM_OIV_305 | 739 | 50.10 | 31.28 | 0.00 | 25.00 | 80.00 | 100.00 |
| LEAF_ERINOSE_PC | 739 | 0.64 | 0.92 | 0.00 | 0.00 | 1.00 | 3.00 |
| MORPHO_OIV_006 | 738 | 3.27 | 1.60 | 1.00 | 2.00 | 5.00 | 9.00 |
| MORPHO_OIV_152 | 674 | 1.92 | 0.47 | 1.00 | 2.00 | 2.00 | 3.00 |
| FERT_GLOB | 739 | 5.91 | 3.48 | 0.00 | 3.00 | 8.00 | 19.00 |
| FERT_PRIM | 739 | 1.20 | 0.70 | 0.00 | 0.60 | 1.80 | 3.80 |
| FERT_SPURS | 701 | 0.57 | 1.03 | 0.00 | 0.00 | 1.00 | 6.00 |
| SHOOT_L | 739 | 67.01 | 24.80 | 0.00 | 48.50 | 82.20 | 174.20 |
| INT_NB | 739 | 2.47 | 2.81 | 0.00 | 0.00 | 3.80 | 48.50 |
| SHOOT_II_FREQ_0 | 739 | 0.98 | 1.38 | 0.00 | 0.00 | 2.00 | 5.00 |
| INT_L_45 | 739 | 5.16 | 4.77 | 0.00 | 2.00 | 7.18 | 35.90 |
| INT_L | 739 | 5.26 | 1.33 | 0.00 | 4.40 | 6.00 | 11.50 |
| MORPHO_OIV_204 | 673 | 2.25 | 0.75 | 1.00 | 2.00 | 3.00 | 3.00 |
| YIELD_PLANT | 674 | 1.46 | 1.18 | 0.02 | 0.69 | 1.92 | 9.42 |
| NB_CLUST_PLANT | 739 | 7.46 | 4.59 | 0.00 | 4.00 | 10.00 | 25.00 |
| SCLUST_W_G | 674 | 175.72 | 97.45 | 1.80 | 110.08 | 220.07 | 735.00 |
| MORPHO_OIV_240 | 672 | 1.91 | 0.56 | 0.50 | 1.50 | 2.30 | 3.50 |
| SBER_W_g | 674 | 3.04 | 1.23 | 0.41 | 2.06 | 3.87 | 7.47 |
| MUST_Y_PLANT | 674 | 649.94 | 636.88 | 12.00 | 275.00 | 860.00 | 7070.00 |
| MUST_ALC_C | 673 | 11.01 | 2.36 | 2.51 | 9.94 | 12.55 | 17.10 |
| MUST_TA_H2SO4 | 673 | 4.96 | 1.91 | 1.46 | 3.42 | 6.20 | 16.20 |
| MUST_pH | 660 | 3.15 | 0.19 | 2.64 | 3.02 | 3.26 | 4.02 |
| MUST_TAR_H2SO4 | 661 | 6.19 | 1.67 | 0.71 | 5.10 | 7.40 | 11.00 |
| MUST_MAL_gH2SO4l | 661 | 3.04 | 2.18 | 0.00 | 1.46 | 4.10 | 16.70 |
| MUST_K | 661 | 1277.36 | 244.67 | 788.00 | 1081.00 | 1428.00 | 2156.00 |
| MUST_N_AA&NH4 | 657 | 177.48 | 65.86 | 27.00 | 133.00 | 216.00 | 557.00 |
| VIGOUR_VISUAL | 738 | 7.42 | 2.16 | 1.00 | 7.00 | 9.00 | 9.00 |

Table S3 Summary of variables for raw phenotypic data

| EDGARR | | | | | | | | |
|---|---|---|---|---|---|---|---|---|
| trait | missing.dat | ngeno | fixed.effects | random.effects | var.geno | var.resid | var.tot | H2 |
| HARVEST_DOY | 12 | 203 | year | geno, NameCross | 110.02 | 400.03 | 557.32 | 0.438 |
| NB_CLUST_PLANT | 25 | 196 | | geno, row, col | 2.78 | 50.82 | 58.13 | 0.081 |
| BER_GLUC_HPLC_mgber | 61 | 127 | year | geno, row, col | 190.47 | 3,836.27 | 4,557.25 | 0.058 |
| BER_GLUC_HPLC_g | 61 | 127 | year | geno, row, col | 0.21 | 68.08 | 111.78 | 0.003 |
| BER_FRUC_HPLC_mgber | 61 | 127 | year | geno, row, col | 0.04 | 3,692.72 | 4,362.25 | 0.000 |
| BER_TART_HPLC_mgber | 61 | 127 | year | geno, row, col | 2.87 | 42.78 | 49.21 | 0.080 |
| BER_TART_HPLC_g | 61 | 127 | year | geno, row, col | 0.51 | 1.54 | 2.64 | 0.251 |
| BER_MAL_HPLC_mgber | 61 | 127 | year | geno | 74.12 | 105.14 | 179.26 | 0.498 |
| BER_MAL_HPLC | 61 | 127 | year | geno | 4.18 | 2.72 | 6.91 | 0.684 |
| BER_SHIK_HPLC_mgber | 61 | 126 | year | geno | 0.00 | 0.00 | 0.01 | 0.658 |
| BER_SHIK_HPLC_g | 61 | 126 | year | geno, NameCross | 0.00 | 0.00 | 0.00 | 0.759 |
| SBER_W_g | 41 | 168 | | geno, NameCross | 3,684.50 | 11,030.34 | 16,197.25 | 0.427 |
| BER_ALC_C | 41 | 168 | | geno, row, col | 0.13 | 0.85 | 1.01 | 0.195 |
| BER_TA_g | 42 | 166 | year | geno, row, col | 0.61 | 1.53 | 2.14 | 0.387 |
| BERRY_pH | 43 | 166 | year | geno, row, col | 0.01 | 0.04 | 0.05 | 0.342 |
| BERRY_ANTHO_meqMV | 49 | 150 | | geno, NameCross | 1,181.89 | 3,698.13 | 6,592.34 | 0.549 |
| BER_LIGHT | 41 | 169 | year | geno, NameCross | 58.39 | 154.87 | 289.40 | 0.581 |
| BER_RED | 40 | 169 | | geno, NameCross | 61.87 | 170.49 | 396.58 | 0.681 |
| BER_YELLOW | 40 | 169 | year | geno, row, col | 5.50 | 72.92 | 89.81 | 0.095 |
| OD_420 | 41 | 169 | year | geno | 0.03 | 0.12 | 0.15 | 0.274 |
| OD_520 | 41 | 169 | | geno, NameCross | 0.14 | 0.29 | 0.55 | 0.598 |
| OD_620 | 41 | 169 | year | geno, row, col | 0.00 | 0.02 | 0.03 | 0.255 |
| BER_COLOR_INT | 41 | 169 | year | geno, NameCross | 0.32 | 0.93 | 1.50 | 0.492 |
| BER_TINT | 40 | 169 | | geno, NameCross | 0.08 | 0.12 | 0.31 | 0.718 |
| BER_PI420 | 48 | 151 | year | geno, row, col | 0.02 | 0.07 | 0.09 | 0.271 |
| BER_PI520 | 48 | 151 | year | geno, row, col | 0.01 | 0.02 | 0.03 | 0.278 |
| BER_TPI | 49 | 152 | | geno, NameCross | 5.38 | 14.23 | 22.78 | 0.481 |
| BER_HYDROCIN_mgl | 49 | 152 | year | geno, NameCross | 2.95 | 5.99 | 12.66 | 0.632 |
| BER_TOTAL_ANTHO | 48 | 152 | | geno, NameCross | 3.27 | 9.81 | 17.00 | 0.531 |

| Martell | | | | | | | | | | | |
|---|---|---|---|---|---|---|---|---|---|---|---|
| trait | missing.dat | ngeno | semi.quanti | fixed.effects | random.effects | var.geno | var.cross | var.cross.geno | var.resid | var.tot | H2 |
| DBT_RELATIVE | 0 | 358 | FALSE | year | geno | 16.76 | NA | NA | 32.23 | 49.00 | 0.518 |
| FLO_RELATIVE | 10 | 349 | FALSE | year | geno, Position | 1.12 | NA | NA | 7.94 | 9.47 | 0.213 |
| VER_RELATIVE | 11 | 344 | FALSE | year | geno, Cross | 25.23 | 2.90 | 0.10 | 47.17 | 84.21 | 0.601 |
| WOODM_OIV_305 | 0 | 358 | FALSE | year | geno | 27.10 | NA | NA | 509.85 | 536.95 | 0.099 |
| ERINO_LEAF_PC | 0 | 358 | FALSE | year | geno | 0.05 | NA | NA | 0.49 | 0.55 | 0.184 |
| MORPHO_OIV_006 | 0 | 358 | TRUE | year | geno, Cross | 0.26 | 0.03 | 0.09 | 1.59 | 1.90 | 0.287 |
| MORPHO_OIV_152 | 9 | 345 | TRUE | year | geno | 0.03 | NA | NA | 0.18 | 0.21 | 0.277 |
| FERT_GLOB | 0 | 358 | FALSE | year | geno, Cross | 1.36 | 5.29 | 0.80 | 8.22 | 12.48 | 0.517 |
| FERT_PRIM | 0 | 358 | FALSE | year | geno, Cross | 0.06 | 1,925.34 | 1.00 | 0.32 | 0.51 | 0.537 |
| FERT_COURSON | 5 | 358 | FALSE | year | geno | 0.20 | NA | NA | 0.76 | 0.97 | 0.344 |
| SHOOT_L | 0 | 358 | FALSE | | geno, Cross, Rang, Mode_de_conduite | 132.72 | 0.16 | 0.00 | 330.83 | 634.33 | 0.490 |
| SHOOT_II_NB | 0 | 358 | FALSE | year | geno | 0.64 | NA | NA | 3.55 | 4.19 | 0.270 |
| SHOOT_II_FREQ_0 | 0 | 358 | FALSE | year | geno | 0.00 | NA | NA | 0.80 | 0.80 | 0.002 |
| SHOOT_II_L | 0 | 358 | FALSE | | geno | 2.30 | NA | NA | 20.52 | 22.81 | 0.188 |
| INT_L | 0 | 358 | FALSE | | geno, Rang, Mode_de_conduite | 0.20 | NA | NA | 1.43 | 1.81 | 0.224 |
| MORPHO_OIV_204 | 9 | 348 | TRUE | | geno, Cross, Rang, Position | 0.67 | 8,304.02 | 1.00 | 0.02 | 0.94 | 0.987 |
| YIELD_PLANT | 9 | 347 | FALSE | year | geno, Cross, Position | 0.26 | 114.99 | 1.00 | 0.85 | 1.34 | 0.514 |
| NB_CLUST_PLANT | 0 | 358 | FALSE | year | geno, Cross | 2.78 | 11.81 | 0.81 | 14.22 | 22.28 | 0.539 |
| SCLUST_W_G | 9 | 347 | FALSE | year | geno, Cross | 2,741.05 | 0.05 | 0.00 | 4,912.42 | 9,578.82 | 0.649 |
| MORPHO_OIV_240 | 9 | 347 | FALSE | | geno, Cross | 0.08 | 2.90 | 0.97 | 0.19 | 0.32 | 0.570 |
| SBER_W_G | 9 | 347 | FALSE | year | geno, Cross, RUN_RPV | 0.15 | 0.12 | 0.45 | 0.47 | 0.83 | 0.558 |
| MUST_PLANT | 9 | 347 | FALSE | year | geno | 102,540.99 | NA | NA | 268,445.69 | 370,986.67 | 0.426 |
| MUST_TAVP | 9 | 347 | FALSE | | geno | 1.40 | NA | NA | 4.17 | 5.57 | 0.395 |
| MUST_TA | 9 | 347 | FALSE | year | geno, Cross, RPV3, Rang | 0.57 | 0.20 | 0.26 | 0.76 | 1.66 | 0.669 |
| MUST_pH | 11 | 346 | FALSE | year | geno, Cross, Position | 0.00 | 5.29 | 1.00 | 0.01 | 0.03 | 0.681 |
| MUST_TARTRIQUE | 11 | 347 | FALSE | year | geno, Cross, Rang | 0.12 | 1,925.34 | 1.00 | 1.07 | 1.55 | 0.279 |
| MUST_MALIQUE | 11 | 347 | FALSE | year | geno, Cross, RPV3 | 0.93 | 0.05 | 0.05 | 1.12 | 2.84 | 0.725 |
| POTASSIUM | 11 | 347 | FALSE | year | geno, Cross, Rang, Position | 5,927.52 | 0.16 | 0.00 | 25,092.09 | 46,161.80 | 0.519 |
| MUST_NASS | 11 | 344 | FALSE | year | geno, Cross, Rang | 1,189.98 | 0.21 | 0.00 | 2,212.11 | 4,000.39 | 0.530 |
| VIGOUR_VISUAL | 0 | 358 | TRUE | year | geno | 0.25 | NA | NA | 2.73 | 2.98 | 0.158 |

Table S4 Table of fitting information

| POPULATION | EDGARR | MARTELL |
|---|---|---|
| **HALF-DIALLEL** | 3943 SNPs | 2804 SNPs |
| **DIVERSITY PANEL** | 6301 SNPs | 4676 SNPs |
| **HALF-DIALLEL + DIVERSITY PANEL** | 3707 SNPs | 2650 SNPs |

Table S5 number of common SNP markers between populations

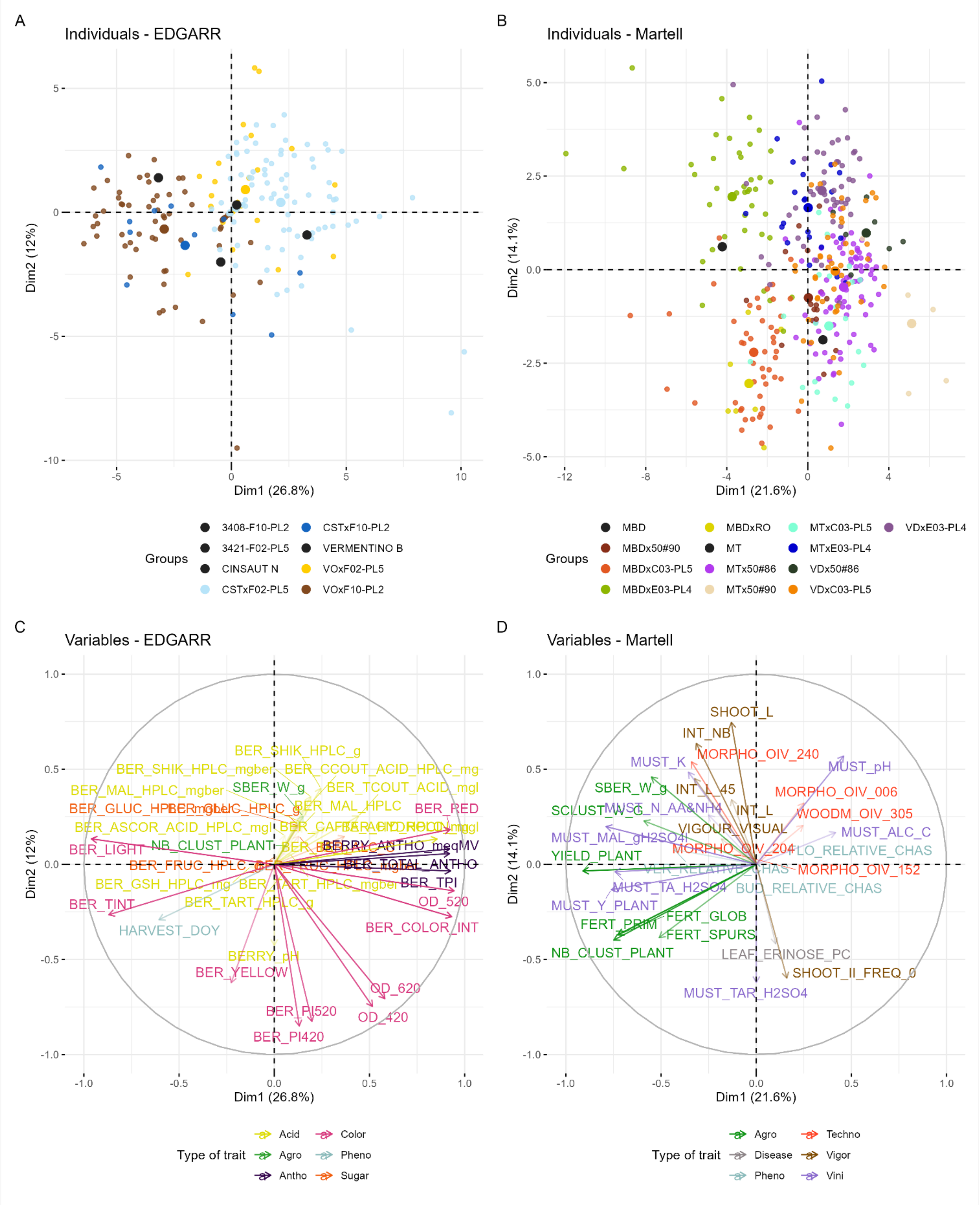

Figure S6 Phenotypic structure, results of PCA analysis. A & B: plot of variable contributions for the first two axes for EDGARR and Martell populations, respectively, C & D: projection of individuals for EDGARR and Martell populations, respectively.

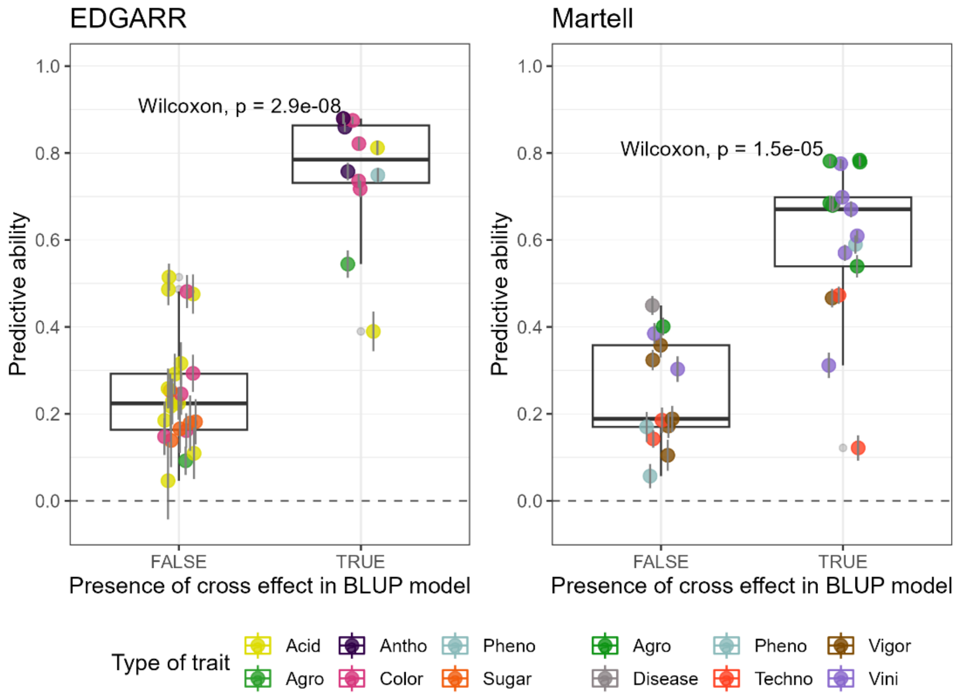

Figure S7 Effect of the presence of cross effect in the mixed model on predictive ability for both populations, for the best method between GBLUP and LASSO. Wilcoxon test was performed between the groups, and a p-value of the test is displayed in the plot.

A. EDGARR

| TRAIT | DIRECTION | WEIGHT |
|---|---|---|
| **BER_TART_HPLC_G** | 9 | 0.49 |
| **BER_LIGHT** | Max | 0.82 |
| **BER_COLOR_INT** | Min | 0.72 |
| **HARVEST_DOY** | Max | 0.75 |
| **BER_MAL_HPLC** | 13 | 0.32 |
| **BER_TA_G** | 7 | 0.29 |
| **BERRY_PH** | 3.5 | 0.26 |
| **BER_ALC_C** | 11 | 0.25 |
| **BER_TPI** | Min | 0.76 |
| **NB_CLUST_PLANT** | Max | 0.092 |
| **SBER_W_G** | Max | 0.54 |

B. Martell

| TRAIT | DIRECTION | WEIGHT |
|---|---|---|
| **MORPHO_OIV_240** | 2 | 0.5 |
| **FERT_GLOB** | Max | 0.5 |
| **FERT_PRIM** | Max | 0.5 |
| **YIELD_PLANT** | Max | 1 |
| **MUST_Y_PLANT** | max | 1 |
| **MUST_MAL_H2SO4** | 3 | 0.5 |
| **MUST_TAR_H2SO4** | Max | 1 |
| **MUST_K** | Min | 0.5 |
| **SCLUST_W_G** | Max | 0.5 |
| **SBER_W_G** | Max | 0.5 |
| **MUST_TA_H2SO4** | Max | 0.5 |
| **MUST_PH** | Min | 0.5 |

Table S8 Ideotype definition, A: for EDGARR, B: for Martell population. The direction column indicates the selection direction, if a numeric value is displayed, an optimum value is sought for this trait.

| EDGARR | | | | | | | |
|---|---|---|---|---|---|---|---|
| VAR | Factor | Xo | Xs | SD | SDperc | sense | goal |
| BER_TART_HPLC_g | FA1 | 1.78 | 2.08 | 0.30 | 16.93 | decrease | 0 |
| BER_LIGHT | FA1 | 68.59 | 70.59 | 2.00 | 2.92 | increase | 100 |
| BER_COLOR_INT | FA1 | 1.65 | 1.53 | −0.13 | −7.69 | decrease | 100 |
| HARVEST_DOY | FA1 | 258.46 | 274.62 | 16.17 | 6.26 | increase | 100 |
| BER_MAL_HPLC | FA2 | 2.19 | 3.04 | 0.85 | 38.91 | decrease | 0 |
| BER_TA_g | FA2 | 1.73 | 2.57 | 0.84 | 48.47 | decrease | 0 |
| BERRY_pH | FA2 | 0.17 | 0.27 | 0.10 | 60.32 | decrease | 0 |
| BER_ALC_C | FA3 | 0.60 | 0.46 | −0.14 | −23.32 | decrease | 100 |
| BER_TPI | FA3 | 11.75 | 8.48 | −3.27 | −27.82 | decrease | 100 |
| NB_CLUST_PLANT | FA3 | 9.86 | 9.30 | −0.56 | −5.64 | increase | 0 |
| SBER_W_g | FA3 | 368.31 | 379.24 | 10.93 | 2.97 | increase | 100 |

| Martell | | | | | | | |
|---|---|---|---|---|---|---|---|
| VAR | Factor | Xo | Xs | SD | SDperc | sense | goal |
| MUST_TA_H2SO4 | FA1 | 6.93 | 7.55 | 0.61 | 8.85 | increase | 100 |
| MUST_pH | FA1 | 3.01 | 2.94 | −0.07 | −2.36 | decrease | 100 |
| YIELD_PLANT | FA1 | 1.37 | 1.65 | 0.28 | 20.30 | increase | 100 |
| FERT_PRIM | FA1 | 1.09 | 1.52 | 0.43 | 39.50 | increase | 100 |
| FERT_GLOB | FA1 | 5.34 | 7.43 | 2.08 | 39.01 | increase | 100 |
| SCLUST_W_G | FA2 | 169.24 | 140.40 | −28.85 | −17.04 | increase | 0 |
| MUST_TAR_H2SO4 | FA2 | 7.44 | 8.41 | 0.97 | 13.05 | increase | 100 |
| SBER_W_g | FA2 | 2.22 | 2.12 | −0.11 | −4.78 | increase | 0 |
| MUST_MAL_gH2SO4l | FA2 | 2.50 | 2.47 | −0.02 | −0.92 | decrease | 100 |
| MORPHO_OIV_240 | FA3 | 0.71 | 0.63 | −0.09 | −12.28 | decrease | 100 |
| MUST_K | FA3 | 1,480.93 | 1,477.97 | −2.95 | −0.20 | decrease | 100 |

Table S9 Summary of MGIDI output for EDGARR and Martell population. Factor: closest factor where each trait belongs to, Xo and Xs: mean of the original and selected population, respectively, SD: standard deviation, SDperc: percentage of standard deviation; sense: direction of the trait for the defined ideotype, goal: 0 or 100 if the selection direction has been respected.

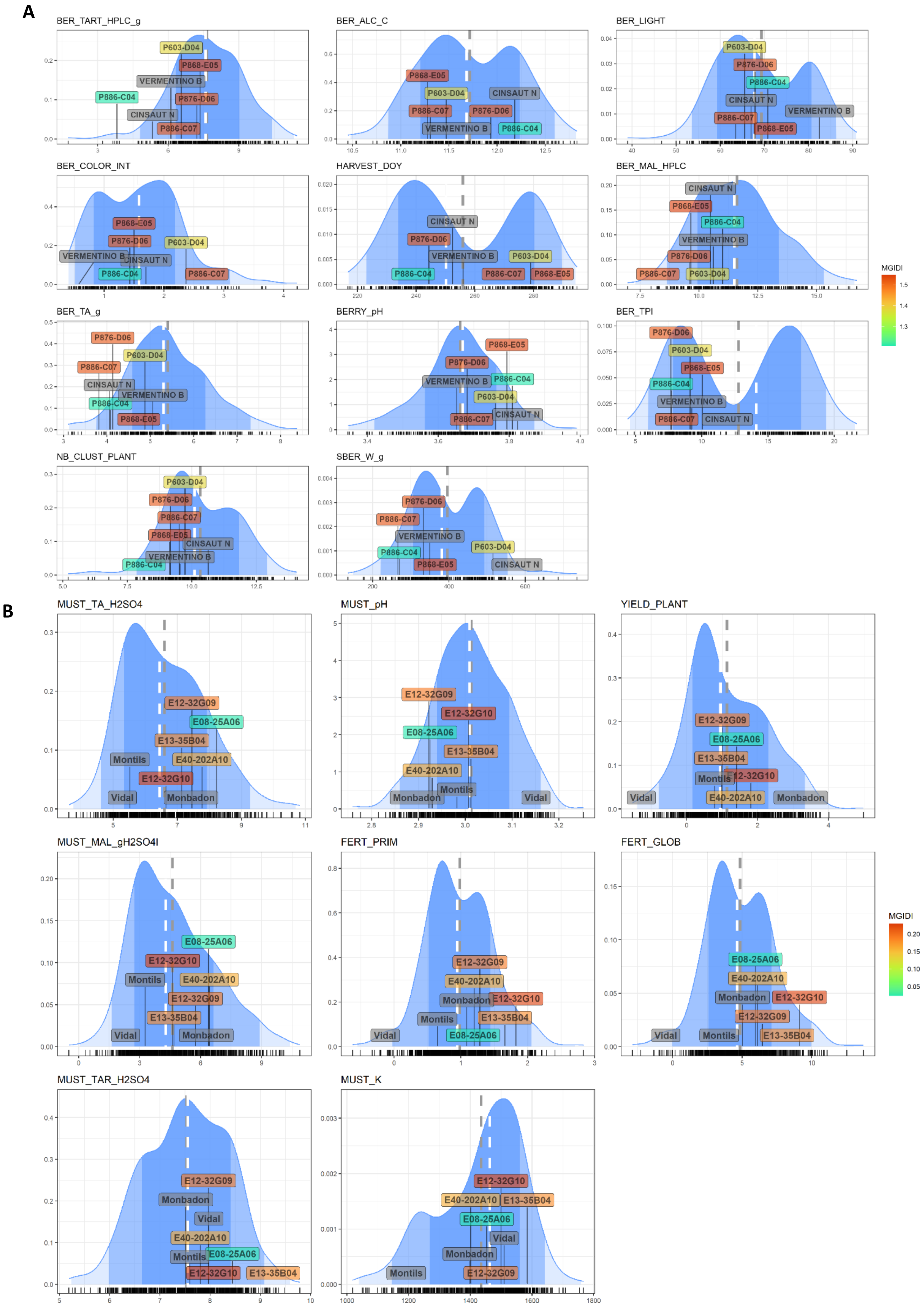

A
BER_TART_HPLC_g
P603-D04
P868-E05
VERMENTINO B
P886-C04
P876-D06
CINSAUT N
P886-C07
BER_ALC_C
P868-E05
P603-D04
CINSAUT N
P886-C07
P876-D06
VERMENTINO B
P886-C04
BER_LIGHT
P603-D04
P876-D06
P886-C04
CINSAUT N
VERMENTINO B
P886-C07
P868-E05
BER_COLOR_INT
P868-E05
P876-D06
P603-D04
VERMENTINO B
CINSAUT N
P886-C04
P886-C07
HARVEST_DOY
CINSAUT N
P876-D06
VERMENTINO B
P603-D04
P886-C04
P886-C07
P868-E05
BER_MAL_HPLC
CINSAUT N
P868-E05
P886-C04
VERMENTINO B
P876-D06
P886-C07
P603-D04
BER_TA_g
P876-D06
P603-D04
P886-C07
CINSAUT N
VERMENTINO B
P886-C04
P868-E05
BERRY_pH
P868-E05
P876-D06
VERMENTINO B
P886-C04
P603-D04
P886-C07
CINSAUT N
BER_TPI
P876-D06
P603-D04
P868-E05
P886-C04
VERMENTINO B
P886-C07
CINSAUT N
NB_CLUST_PLANT
P603-D04
P876-D06
P886-C07
P868-E05
CINSAUT N
VERMENTINO B
P886-C04
SBER_W_g
P876-D06
P886-C07
VERMENTINO B
P603-D04
P886-C04
P868-E05
CINSAUT N
MGIDI
1.5
1.4
1.3
B
MUST_TA_H2SO4
E12-32G09
E08-25A06
E13-35B04
Montils
E40-202A10
E12-32G10
Vidal
Monbadon
MUST_pH
E12-32G09
E12-32G10
E08-25A06
E13-35B04
E40-202A10
Montils
Monbadon
Vidal
YIELD_PLANT
E12-32G09
E08-25A06
E13-35B04
Montils
E12-32G10
Vidal
E40-202A10
Monbadon
MUST_MAL_gH2SO4l
E08-25A06
E12-32G10
Montils
E40-202A10
E12-32G09
E13-35B04
Vidal
Monbadon
FERT_PRIM
E12-32G09
E40-202A10
Monbadon
E12-32G10
Montils
E13-35B04
Vidal
E08-25A06
FERT_GLOB
E08-25A06
E40-202A10
Monbadon
E12-32G10
E12-32G09
Vidal
Montils
E13-35B04
MGIDI
0.20
0.15
0.10
0.05
MUST_TAR_H2SO4
E12-32G09
Monbadon
Vidal
E40-202A10
Montils
E08-25A06
E12-32G10
E13-35B04
MUST_K
E12-32G10
E40-202A10
E13-35B04
E08-25A06
Vidal
Monbadon
Montils
E12-32G09

Figure S10 Distribution of genotypic values in the validation population and position of the first five individuals and the emblematic parents. A: EDGARR population, B: Martell population.

| EDGARR | | | |
|---|---|---|---|
| Genotype | Cross | color | MGIDI |
| P886-C04 | Vermentino x COL48-59 | white | 1.24 |
| P603-D04 | Cinsaut x F10-PL2 | red | 1.29 |
| P868-E05 | Vermentino x F10-PL2 | red | 1.39 |
| P249-F10 | Cinsaut x F10-PL2 | red | 1.43 |
| P865-D07 | Vermentino x F10-PL2 | red | 1.48 |
| P876-D06 | Vermentino x COL48-46 | white | 1.51 |
| P624-A09 | Vermentino x F10-PL2 | red | 1.52 |
| P869-F04 | Vermentino x F10-PL2 | red | 1.59 |
| P886-C07 | Vermentino x COL48-59 | red | 1.65 |
| P596-A09 | Cinsaut x F10-PL2 | red | 1.79 |
| P867-D05 | Vermentino x F10-PL2 | red | 1.79 |
| P862-E05-D | Vermentino x F10-PL2 | red | 1.87 |
| P869-B07 | Vermentino x F10-PL2 | white | 1.87 |
| P875-D03 | Vermentino x COL48-46 | red | 1.89 |
| P865-B05 | Vermentino x F10-PL2 | white | 1.92 |

| Martell | | |
|---|---|---|
| Genotype | Cross | MGIDI |
| E12-32G10 | Monbadon x C03-PL5 | 0.19 |
| E01-2B03 | Monbadon x Rayon d'Or | 0.27 |
| E12-32G09 | Monbadon x C03-PL5 | 0.35 |
| E01-1C13 | Monbadon x Rayon d'Or | 0.42 |
| E09-26G04 | Monbadon x C03-PL5 | 0.43 |
| E59-254B06 | C03-PL5 x Vidal | 0.49 |
| E10-29D10 | Monbadon x C03-PL5 | 0.56 |
| E11-30C02 | Monbadon x C03-PL5 | 0.57 |
| E08-25A06 | Monbadon x C03-PL5 | 0.58 |
| E11-32A03 | Monbadon x C03-PL5 | 0.63 |
| E11-31A10 | Monbadon x C03-PL5 | 0.67 |
| E08-25A03 | Monbadon x C03-PL5 | 0.69 |
| E13-34H07 | Monbadon x C03-PL5 | 0.70 |
| E59-253H12 | C03-PL5 x Vidal | 0.70 |
| E09-27D01 | Monbadon x C03-PL5 | 0.73 |

Table S11 Table of the 15 best individuals for both populations. MGIDI: multi-trait genotype-ideotype distance index. For the EDGARR population, the LASSO method predicted berry color based on genomic prediction.

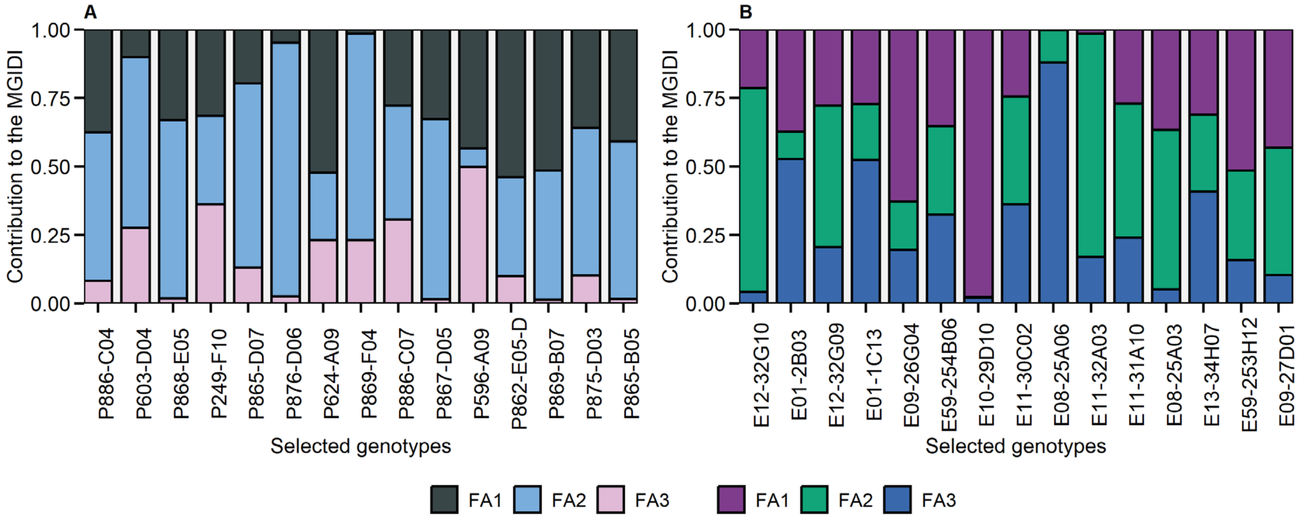

Figure 12 Strength and weakness of selected genotypes.

The Y-axis represents the contribution of each factor to the distance to the ideotype. The higher the contribution, the less the genotype is close to the ideotype. Traits contributing to each factor are displayed in Figure S9. A: EDGARR population, B: Martell population
